# The Tangled History and Taxonomy of an Iconic Chorus Frog Complex Clarified using Genomic Analyses

**DOI:** 10.64898/2026.03.09.710633

**Authors:** Chris X. McDaniels, Nicole Povelikin, Mysia Dye, Michelle L. Kortyna, Robert C. Jadin, Sarah A. Orlofske, Gabriela Parra-Olea, Alan R. Lemmon, Emily Moriarty Lemmon, Lisa N. Barrow

**Author notes:** Correspondence to be sent to: Department of Biology, University of New Mexico, MSC03-2020, 219 Yale Blvd NE, Albuquerque, NM 87131, USA.

## Abstract

Species represent a fundamental unit of biodiversity in evolutionary biology, but the nature of the speciation continuum and inadequate sampling of organisms with broad distributions provide substantial challenges to species delimitation. The Pacific Treefrog complex (*Pseudacris regilla sensu lato*) is an iconic but systematically poorly understood group of chorus frogs inhabiting a vast portion of western North America. Current studies tentatively recognize three species in this complex (*P. hypochondriaca*, *P. regilla*, *P. sierra*), but disagreement remains among morphological, mitochondrial, and nuclear genetic data. In this study, we used thorough geographic sampling and thousands of nuclear loci, along with an integrative, multi-method approach to clarify the phylogenetic relationships and divergence history of *P. regilla s.l.* lineages and recommend a new taxonomic arrangement for the group. *Pseudacris regilla* and *P. sierra* fall firmly within the “gray zone” of speciation, composing a combined “north” lineage. Based on the degree of congruence in inferences from our analyses and evidence for isolating mechanisms, we propose a two species taxonomy for this complex, recognizing the “north” lineage as *P. regilla* and retaining *P. hypochondriaca* as a species. Our study shows how extensive geographic sampling, high-throughput sequencing, and multiple analytical approaches can resolve systematic uncertainties in challenging species complexes.

---

Understanding species diversity is an essential goal in systematic biology, as species represent definable units used in communicating about and studying biodiversity. Despite this importance, delimiting species remains challenging because of the nature of speciation and the limitations inherent in sampling organisms with broad geographic distributions. Speciation is a continuum that often has fuzzy edges; however, species delimitation requires applying discrete categories to a frequently non-instantaneous process. Challenges then arise in trying to define species, particularly when species definitions come into conflict for populations that are less discrete, referred to as the “gray zone” of speciation (de Queiroz 2007). Efforts to reconcile alternative species concepts with the reality of a speciation continuum have led to proposals for a unified species concept (de Queiroz 1998; de Queiroz 2007). This concept only requires species to be “separately evolving metapopulation lineages”, recognizing that additional properties that may develop during the speciation process, such as reproductive isolation or ecological distinctness, can represent evidence supporting species status, but are not explicitly required nor a guarantee that the lineages are different species (de Queiroz 1998; de Queiroz 2007). Ideally, multiple lines of evidence can be gathered that corroborate species boundaries.

Gathering evidence to delimit species is especially challenging for species with broad geographic and environmental distributions. Such species can exhibit strong patterns of Isolation-by-Distance (IBD), in which population genetic differentiation is correlated with geographic distance (Wright 1943). Sampling strategies that ignore intermediate populations between geographical or environmental extremes may erroneously infer discrete population structure (Frantz et al. 2009; Perez et al. 2018) that could be interpreted as interspecific variation (Mason et al. 2020; Chambers et al. 2023). Alternatively, even when geographic sampling is sufficient, the type and amount of data used can affect species delimitation. Discordance among morphology, mitochondrial DNA, and nuclear DNA is common (Rubinoff and Holland 2005; Toews and Brelsford 2012; Bonnet et al. 2017; Ruiz-García et al. 2018), and sampling few genes may not capture the full evolutionary history of a study system because of stochastic error and gene tree discordance (Rokas et al. 2003; Degnan and Rosenberg 2009). Genomic datasets spanning hundreds or thousands of loci have offered new opportunities to revisit previous studies, enabling better assessments of evolutionary history. Recent approaches have successfully used large genomic datasets and multiple lines of evidence to delimit species in the face of the challenges inherent to the speciation process (Chambers et al. 2025b; Waldron et al. 2025).

The Pacific Treefrog complex (*Pseudacris regilla sensu lato*) is an excellent study system for tackling challenging problems in species delimitation. This complex occurs across much of the western coast of North America, from southern Baja California Sur into British Columbia and introduced populations in southeastern Alaska, throughout contrasting environments such as the Mojave Desert and the Pacific temperate rainforests, and from elevations at sea level to over 3,000 meters in the Sierra Nevada mountain range (Stebbins and McGinnis 2018; Spencer 2025). This system has been studied for decades across evolutionary ecology (Morey 1990; Watkins 1996), disease ecology (Orlofske et al. 2012), physiology (Pfab et al. 2020; Montana et al. 2023), and behavior (Pearl et al. 2003; Nelson et al. 2017). In addition to its continuous use in research, *P. regilla sensu lato* (*s.l.)* is one of the most well-recognized frogs in North America, and perhaps the world, because its familiar “ribbit” call is commonly heard in the background of Hollywood films (Elliot et al. 2009; Stebbins and McGinnis 2018). Despite the long history of human interactions with these frogs and their status as a Hollywood icon, it is still unclear whether this “Hollywood frog” represents one cosmopolitan species, three distinct species, or a different number of species.

The history of systematic work in the *Pseudacris regilla s.l.* complex has spanned the use of multiple data types producing conflicting results. Early work using morphological characteristics, including coloration, markings, and body measurements, recognized up to ten subspecies in the *P. regilla s.l.* complex (Jameson et al. 1966) (Fig. 1a).

**FIGURE 1.**
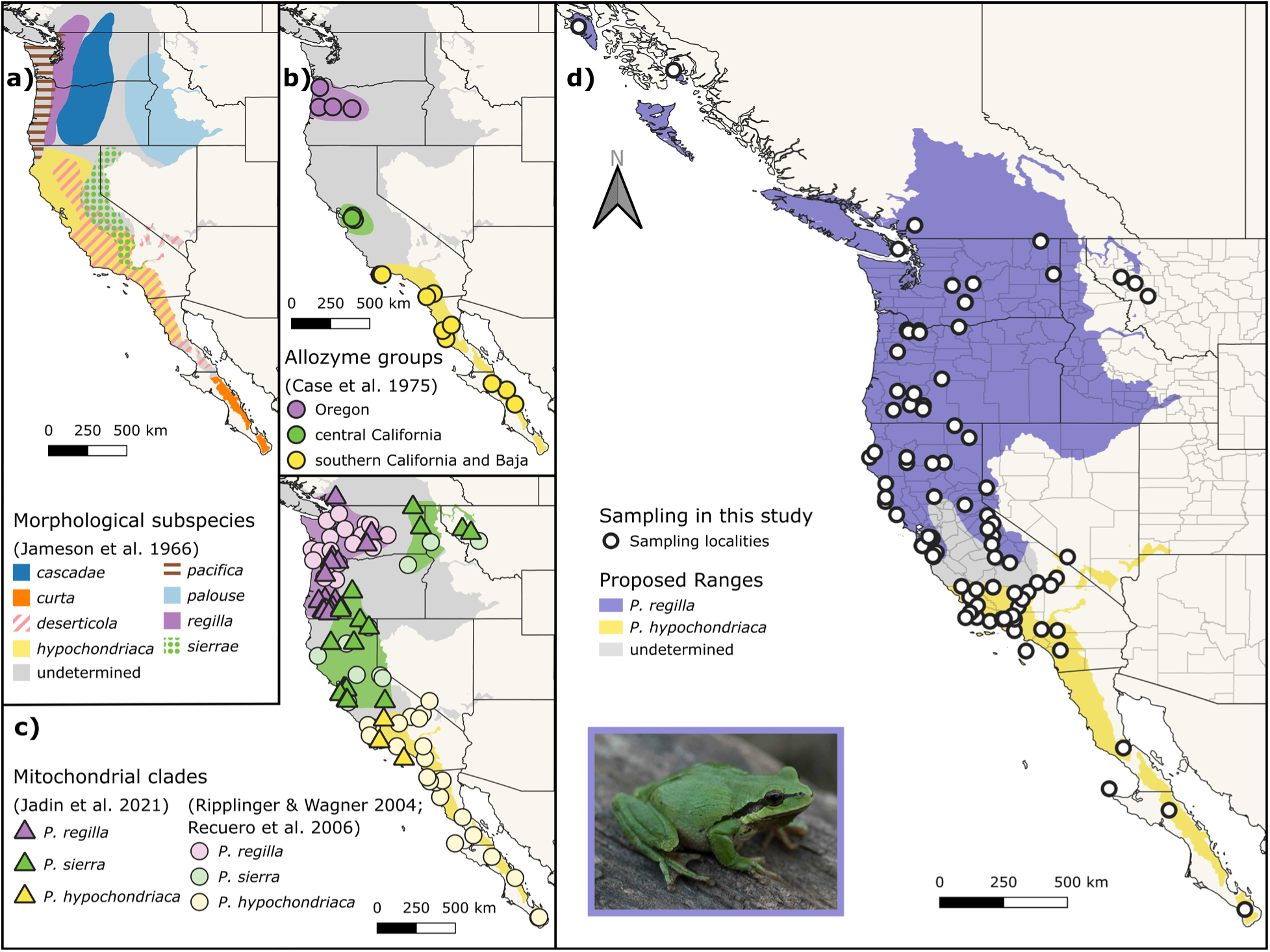
Taxonomic arrangements and geographic ranges of the *Pseudacris regilla sensu lato (s.l.)* complex based on different data types. Ranges are approximate and drawn based on maps or sampling localities in the source publications. Light gray “undetermined” coloring represents unsampled areas that could not be assigned to a taxon. Symbols represent sampling localities. a) Morphological subspecies described in Jameson et al. (1966). b) Allozyme groups described by Case et al. (1975). c) Mitochondrial clades described by Ripplinger and Wagner (2004), Recuero et al. (2006), and Jadin et al. (2021). See note about identification and locality changes in Supplementary Materials. d) Nuclear genomic loci used in this study with proposed ranges and taxonomic designations. Photograph of *P. regilla* by Moses Michelsohn, used with permission. Range map for the *P. regilla s.l.* complex modified from shapefiles courtesy of Travis W. Taggart (cnah.org). Country and state shapefile from the Commission for Environmental Cooperation (including Natural Resources Canada, Instituto Nacional de Estadística y Geografía, and National Atlas of the United States). County shapefile from the National Atlas of the United States. Alt text: Four maps depicting geographic ranges for the *Pseudacris regilla sensu lato* complex based on data type. Colors and patterns are used to represent different taxa ranges.

**FIGURE 2.**
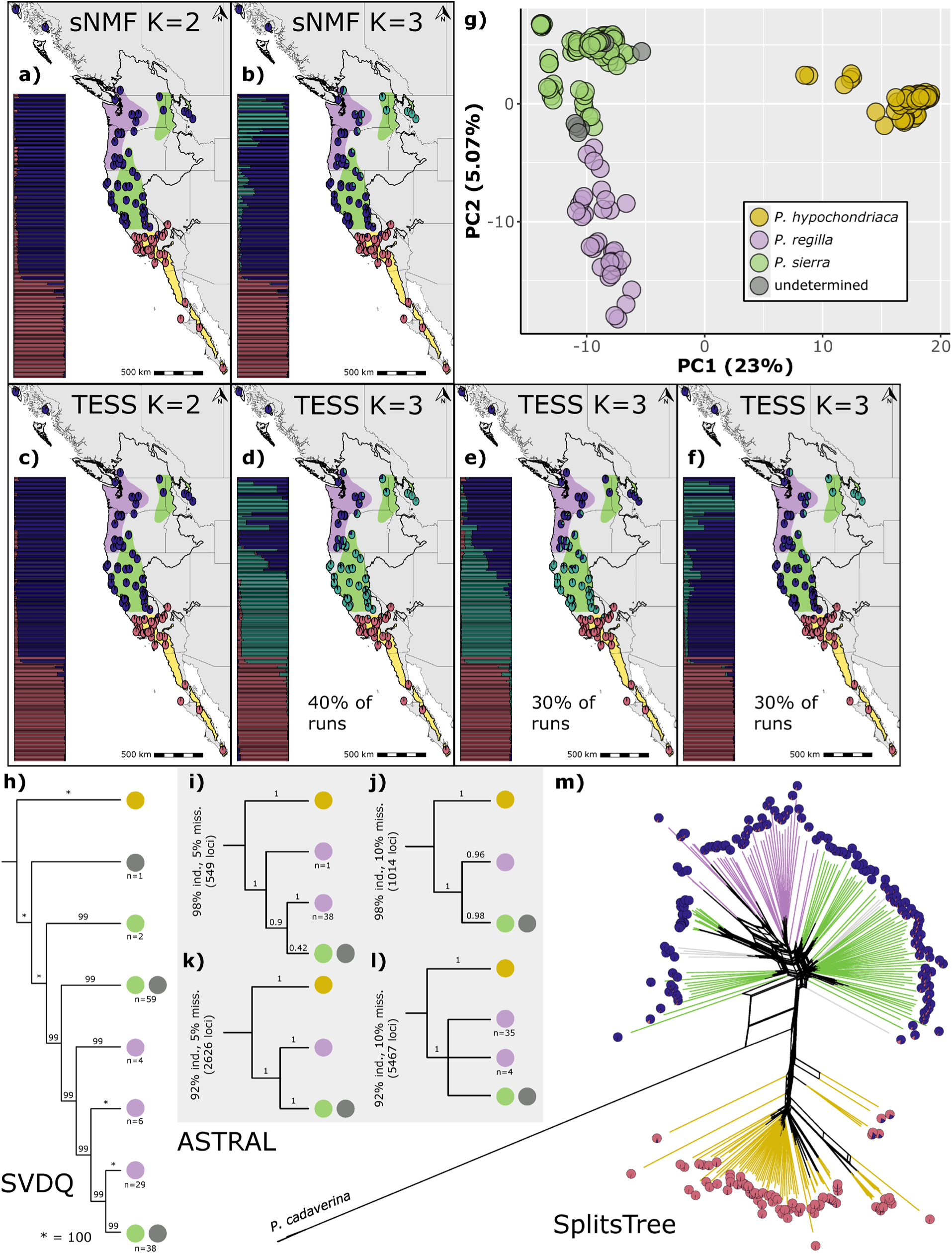
Population structure and phylogenies. a – b) sNMF results for two ancestral population clusters (K = 2) and K = 3, respectively. Each pie chart represents an individual sample and its ancestry proportions. Bar plots show the same ancestry proportions and are sorted by latitude. Colored ranges are based on ranges for *Pseudacris hypochondriaca*, *P. regilla*, and *P. sierra* we approximated from the localities in Jadin et al. (2021). The complete range of *P. regilla sensu lato* is outlined in black. c) TESS3 results for K = 2. d – f) Three alternative TESS3 results for K = 3. g) Principal Component Analysis plot for Principal Components 1 and 2. Samples are colored by species based on Jadin et al. (2021). Samples labeled “undetermined” represent localities that did not fall within the known species ranges inferred from Jadin et al. (2021). h) Simplified SVDQuartets (SVDQ) phylogeny. Bootstrap values are included for each branch. Colored circles at the tips of branches represent species assignments based on Jadin et al. (2021). Number of individuals at a branch tip are shown if that branch does not include all samples assigned to that species (i.e. species is non-monophyletic). i – l) Simplified ASTRAL phylogenies for the four sequence datasets: at least 98% of individuals and no more than 5% missingness, 98% individuals and 10% missingness, 92% individuals and 5% missingness, and 92% individuals and 10% missingness, respectively. m) SplitsTree network with branches colored based on Jadin et al. (2021) and ancestry proportion pie charts on the end of each branch from the TESS3 K = 2 analysis. Alt text: Panels displaying population structure and phylogenetic results. sNMF and TESS3 results are displayed as pie charts on maps.

Later, allozyme analyses identified three groups within *P. regilla s.l.* that corresponded with individuals occurring in Oregon, central California, and southern California and the Baja California peninsula (Case et al. 1975) (Fig. 1b). These three groups were corroborated using cytochrome *b* (cytb) and 12S-16S mitochondrial (mtDNA) genes from across the *P. regilla s.l.* range distribution (Ripplinger and Wagner 2004; Recuero et al. 2006; Jadin et al. 2021) (Fig. 1c). These groups were described as three species by Recuero et al. (2006) and reaffirmed by Jadin et al. (2021): (1) *P. regilla* in the Pacific Northwest, (2) *P. sierra* in central California, Idaho, Montana, eastern Oregon, and eastern Washington, and (3) *P. hypochondriaca* in southern California, southern Nevada, and the Baja California peninsula, with *P. hypochondriaca* and *P. sierra* recognized as sister species, and *P. regilla* sister to that clade. Hereafter, *P. regilla*, unless otherwise noted, refers to the mitochondrial lineage found in the Pacific Northwest. Phylogenetic studies of the *Pseudacris* genus with multi-locus nuclear (nDNA) data have included limited geographic sampling of these three taxa (Barrow et al. 2014; Banker et al. 2020), leaving open questions about the taxonomic and geographic range boundaries within this species complex. Most recently, Vélez and Ingram (2025) found significant differences in reproductive calls between mitochondrial lineages of *P. hypochondriaca* and California populations of *P. sierra*, though no *P. regilla* were included in their study. Currently, the Society for the Study of Amphibians and Reptiles recognizes *P. hypochondriaca*, *P. regilla*, and *P. sierra* (Mendelson III et al. 2025). However, they state, “The final resolution of this taxonomic question awaits nDNA analysis.”

In this study, we used extensive geographic and taxonomic sampling and thousands of nuclear loci to investigate the evolutionary history of the three described species in the *P. regilla s.l.* complex and test the previously generated species hypotheses. Specifically, we asked: (1) What are the phylogenetic relationships within this complex, and how well supported are they? (2) How many species should be delimited within this complex, and where are the species boundaries? (3) What is the divergence history between the lineages within this complex, and is there evidence for historical gene flow or secondary contact? Our detailed sampling and analyses clarify the genomic boundaries in this group and demonstrate the utility of multiple analytical approaches to resolve long-standing taxonomic confusion in broadly distributed species complexes within the gray zone of speciation.

## Materials and Methods

### Sampling, Data Collection, and Data Processing

We obtained three *P. cadaverina* and 227 *P. regilla s.l.* tissue samples from across the range of the *P. regilla s.l* species complex, representing all three putative species and individuals from potential contact zones (Fig. 1d, Supplementary Table S1). Tissues were sourced from previous collections described in Recuero et al. (2006), Barrow et al. (2014), and Jadin et al. (2021) or were requested through loans from the following museum collections: Fort Hays Sternberg Museum of Natural History (FHSM), Museum of Vertebrate Zoology (MVZ), University of Alaska Museum of the North (UAM), and University of Washington Burke Museum (UWBM).

We extracted genomic DNA from muscle or liver tissue using Omega Bio-tek E.Z.N.A. tissue DNA kits (Omega Bio-tek, Norcross, GA), followed by DNA quantification via a Qubit HS dsDNA fluorometric assay (Life Technologies Corporation, Eugene, OR) and visualization of fragment lengths on 2% agarose gels. Using the Fragmentation through Polymerization method (Ignatov et al. 2019), we fragmented 200μg of each extraction into 500–700 base pair (bp) fragments and then performed blunt-end repair and P7 Illumina adapter ligation (Lemmon et al. 2012). We amplified libraries using a tailed primer containing an indexed adapter sequence on the 5’ end, a *Pseudacris*-specific primer on the 3’ end, and a second indexing primer matching to the P7 adapter. Libraries were cleaned, pooled, and sequenced on an Illumina NovaSeq 6000 at the Florida State University College of Medicine Translational Science Lab. We mapped quality-filtered reads to reference sequences from unpublished *P. feriarum* whole-genome sequencing reads and then aligned haplotype sequences across individuals. We discarded haplotype sequences derived from fewer than five reads or paralogous loci and extracted one single nucleotide polymorphism (SNP) per locus using scripts modified from Barrow et al. (2018).

We prepared three different datasets depending on analysis requirements. First, the original SNP dataset included the three *P. cadaverina* samples as an outgroup. Second, we used VCFtools v0.1.17 (Danecek et al. 2011) to create a new VCF file (“ingroup-only”) that excluded the *P. cadaverina* samples and removed monomorphic loci. Third, we created a sequence dataset using Muscle version 5.2.win64 [00ece7c] (Edgar 2022) to align sequences within each locus. Because heterozygous individuals had two sequences representing alternative alleles, we used a custom script that collapsed alleles into one sequence and replaced alternative sites with IUPAC ambiguity codes (available through Dryad). ChatGPT was used to assist with troubleshooting code and writing scripts used in this work. Detailed methods for the following analyses are provided in Supplementary Materials.

### Population Structure and Admixture

We performed several population structure analyses using our ingroup-only SNP dataset. These analyses serve as an initial group discovery step to identify genetic population clusters and the degree of admixture between these populations. In R v4.4.2 (R Core Team 2024), we first ran a Sparse Non-negative Matrix Factorization (sNMF) analysis in the LEA package to infer population genetic structure and estimate admixture coefficients for each individual sample from a genotype matrix (Frichot and François 2015). The snmf function was run with K of 1-15 using 100 repetitions, and entropy was set to TRUE to estimate the cross-entropy criterion for each run. STRUCTURE-like analyses may erroneously infer discrete rather than continuous population structure due to clinal patterns resulting from IBD (Wright 1943; Frantz et al. 2009; Perez et al. 2018). To account for this possibility, we used TESS3 implemented in the algatr R package (Caye et al. 2016; Chambers et al. 2025a), which incorporates spatial statistics into the matrix factorization method used in sNMF. We used the tess_ktest function to perform K selection for K of 1-10. This function selects K based on minimizing the slope between cross-entropy scores, effectively selecting where the cross-entropy scores form an “elbow” or start to plateau. We ran tess_ktest ten times to account for differing outcomes because initial testing showed varying “best” values for K and different clustering patterns. Finally, we ran a Principal Component Analysis (PCA) to visualize population clusters along the two axes explaining the highest proportion of variance in the data. We used the gl.pcoa and gl.pcoa.plot functions in the dartR R package (Gruber et al. 2018; Mijangos et al. 2022) to run the PCA and visualize the results in a biplot.

### Phylogenetic Analyses

We ran several phylogenetic analyses to infer the phylogenetic relationships between the groups discovered by the structure analyses. We used SVDQuartets implemented in PAUP* v4.0a (build 168) to infer a phylogeny from our original SNP dataset (Swofford 2003; Chifman and Kubatko 2014, 2015). SVDQuartets generates sets of quartet splits from single site data to build species trees. We converted our VCF file to NEXUS format using the python script vcf2phylip v2.0 and set --min-samples-locus to 0 to retain all loci (Ortiz 2019). In PAUP*, we defined the three *P. cadaverina* samples as the outgroup and used 10,000,000 random quartets (representing 10.92% of all distinct quartets), 100 bootstrap replicates, and all other settings set to defaults.

We additionally performed phylogenetic analyses using our full sequence data. The final sequence alignments were used to build individual gene trees in IQTree v2.4.0 using 1000 ultrafast bootstraps, and the integrated ModelFinder was used to infer the best substitution model for each locus (Kalyaanamoorthy et al. 2017; Hoang et al. 2018; Minh et al. 2020). The resulting alignments were used as input for inferring species trees under the multi-species coalescent model implemented in ASTRAL v5.7.8 (Zhang et al. 2018). We treated each individual sample as its own taxon. Because some of the sequence alignments contained large gaps or were missing individual samples, we generated four datasets with different thresholds for the percentage of the alignment that contained gaps and the number of missing individuals: (1) 549 loci with at least 98% of individuals (n=214) and up to 5% missing sites, (2) 1014 loci with at least 98% of individuals (n=214) and up to 10% missing sites, (3) 2626 loci with at least 92% of individuals (n=201) and up to 5% missing sites, and (4) 5467 loci with at least 92% of individuals (n=201) and up to 10% missing sites.

We generated a phylogenetic network to visualize any potential non-treelike relationships because our population structure analyses indicated extensive admixture and inconsistent patterns between populations in regions associated with *P. regilla* and *P. sierra*. We converted our VCF file to a distance matrix using the stamppNeisD function in the StAMPP R package and exported it in PHYLIP format using the stamppPhylip function (Pembleton et al. 2013). We used the Neighbor Net algorithm implemented in SplitsTree App v6.4.17 to generate the unrooted phylogenetic network (Bryant and Moulton 2004; Huson and Bryant 2024).

### Species Delimitation

Following the discovery of genetic groups from the structure analyses and the inference of their phylogenetic relationships, we implemented several methods to evaluate these groups from a species delimitation perspective. We created two SuperSOMs (multi-layer Self-Organizing Maps) using the delim-SOM R package (Pyron et al. 2023b). SuperSOMs use unsupervised machine learning to integrate multiple data layers into cluster assignment and species delimitation. SuperSOMs do not rely on *a priori* biological assumptions about the data, and thus simultaneously discover groups and evaluate them rather than evaluating species based on pre-assigned populations. They can detect K = 1, avoid taxonomic over-splitting, and ensure that layer size, such as large SNP matrices, cannot overwhelm other dataset layers.

Our first SuperSOM used our nuclear genomic dataset generated in this study and included alleles, space (latitude, longitude, elevation), and climate (bioclimatic variables) as input layers. We retained only six bioclimatic variables after identifying correlated variables using the corSelect function in the fuzzySim package (Barbosa 2015). We normalized the space and climate data to a scale of 0–1 using the delim-SOM minmax function (Pyron et al. 2023b) and the dplyr mutate function (Wickham et al. 2026) to put them on the same scale as the allele frequency data. We created an output grid using the somgrid function in the delim-SOM package (Pyron et al. 2023b) and then ran a three-layer SuperSOM using the Climate.SOM function (Pyron et al. 2023b) including 100 replicates and 100 steps. We also ran a DNA SOM using only the alleles layer in the DNA.SOM function (Pyron et al. 2023b) to make direct comparisons to our sNMF analysis.

Our second SuperSOM used data from Vélez and Ingram (2025) including alleles (mitochondrial cytochrome *b* sequences), space (latitude, longitude, elevation), climate (bioclimatic variables), and traits (reproductive call properties). We tried coding each haplotype as its own variable in the alleles layer, but there were too many unique haplotypes to produce signal in the results. We therefore treated variations in the sequence alignment as SNPs but acknowledge the potential of introducing pseudo-replication. As input for the traits layer, we chose the mean for each of nine call variables from the locality where each sequence was collected in Vélez and Ingram (2025). We additionally used the latitude, longitude, elevation, and bioclimatic variables provided in Vélez and Ingram (2025). We retained the same six bioclimatic variables as in the first SuperSOM. We normalized all space, climate, and trait variables to a scale of 0–1 as described previously. We ran a four-layer SuperSOM using the Trait.SOM function (Pyron et al. 2023b) with the same input grid, number of replicates, and number of steps as in our first SuperSOM.

To incorporate demographic scenarios involved in lineage formation, we used the species delimitation R package delimitR (Smith and Carstens 2020). delimitR uses fastsimcoal2 simulations to model different combinations of number of species, presence and timing of gene flow, and number of migration edges (Excoffier et al. 2013; Smith and Carstens 2020). The program then uses a random forest (RF) classifier to select which model best reflects the data. We assigned our samples to either “hypo”, “regilla”, or “sierra” based on their positions in the ASTRAL trees, though we note that the evidence for three groups, especially the designations for *P. regilla* and *P. sierra*, are equivocal considering the inconsistent results for K = 3 in the population structure analyses, the differing tree topologies, and the lack of detection for K = 3 by delim-SOM. These assignments were kept to evaluate the previous species hypotheses generated by Recuero et al. (2006) and Jadin et al. (2021) and the demographic history of this group using a model-based method (assignments indicated in Supplementary Table S1). The ingroup-only VCF file was converted to the multi-dimensional site frequency spectrum (mSFS) using easySFS v0.0.1 and downprojected to 76,188,142 (regilla,sierra,hypo) because these values maximized the number of segregating sites (Gutenkunst et al. 2009; Overcast 2023). We changed the first cell in the output file to “0” to remove monomorphic sites. In delimitR, we used the setup_fsc2 function to create nine models (Fig. 3h): one species (Model 1), two species without gene flow (*P. hypochondriaca* and “north”; hereafter, “north” refers to the nuclear genomic clade containing all *P. regilla* and *P. sierra* samples) (Model 2), two species with secondary contact (Model 3), two species with divergence with gene flow (Model 4), three species without gene flow (Model 5), three species with secondary contact between *P. regilla* and *P. sierra* (Model 6), three species with secondary contact between *P. hypochondriaca* and *P. sierra* (Model 7), three species with secondary between *P. regilla* and *P. sierra* and between *P. hypochondriaca* and *P. sierra* (Model 8), and three species with divergence with gene flow between *P. regilla* and *P. sierra* (Model 9).

**FIGURE 3.**
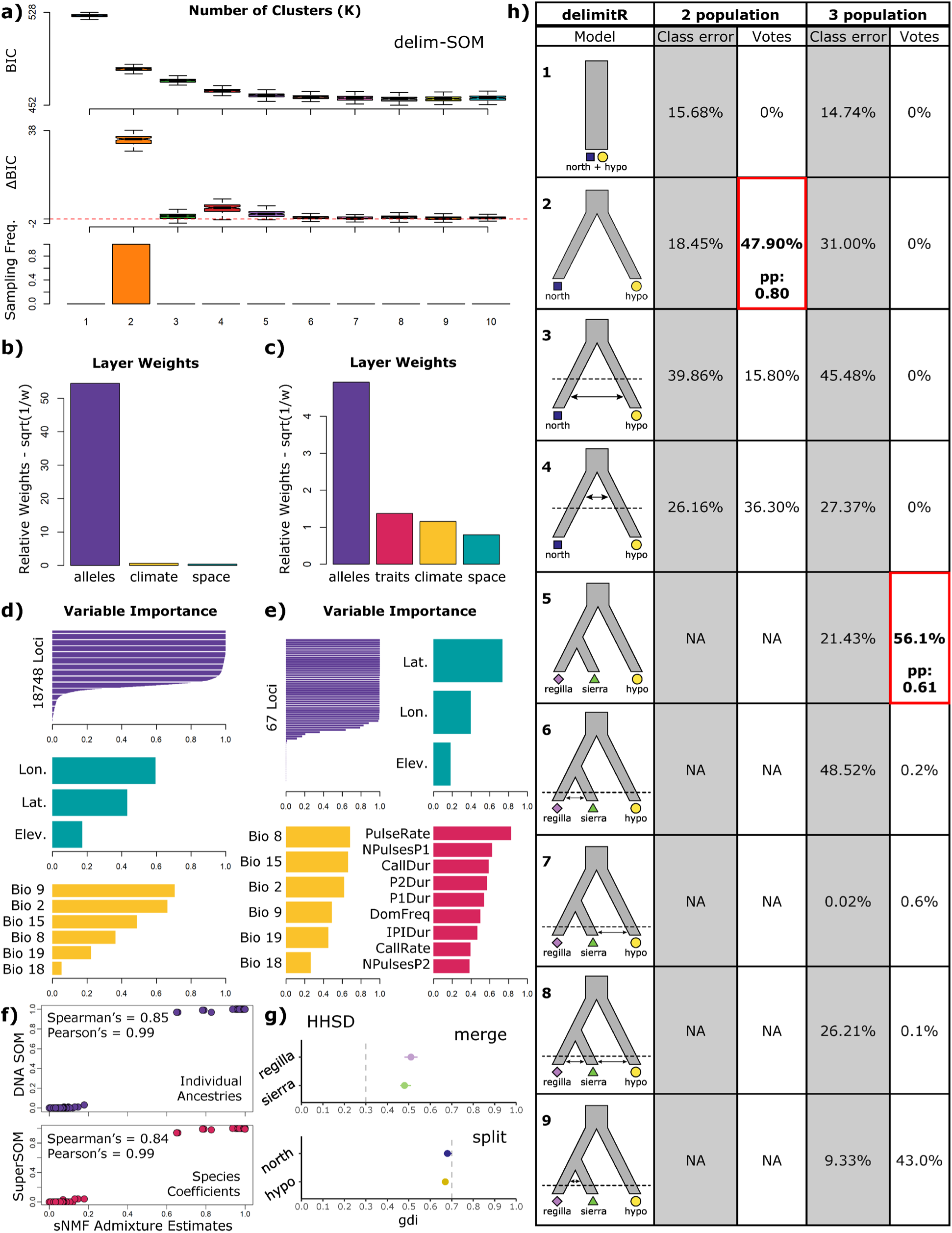
Species delimitation analyses. a) Optimal number of clusters (K) for the three-layer SuperSOM using the nuclear genomic dataset generated in this study. Bayesian Information Criterion, change in Bayesian Information Criterion, and the sampling frequency for each K of 1-10 are reported. b) Layer weights for the three-layer SuperSOM, including alleles, climate, and space layers. c) Layer weights for the four-layer SuperSOM using data from Vélez and Ingram (2025), including alleles, traits, climate, and space layers. d) Variable importance estimates for variables in each layer of the three-layer SuperSOM. For the climate layer, Bio 9 = Mean Temperature of Driest Quarter, Bio 2 = Mean Diurnal Range (Mean of monthly (max temp - min temp)), Bio 15 = Precipitation Seasonality (Coefficient of Variation), Bio 8 = Mean Temperature of Wettest Quarter, Bio 19 = Precipitation of Coldest Quarter, and Bio 18 = Precipitation of Warmest Quarter. e) Variable importance estimates for variables in each layer of the four-layer SuperSOM. For the traits layer, PulseRate = Pulse rate (pulses/s), NPulsesP1 = Number of pulses in Phase 1, Call Dur = Call duration (ms), P2Dur = Phase 2 duration (ms), P1Dur = Phase 1 duration (ms), DomFreq = Dominant frequency (Hz), IPIDur = Interphase interval duration (ms), CallRate = Call rate (calls/s), and NPulsesP2 = Number of pulses in Phase 2. f) Comparisons between sNMF ancestry coefficients and the DNA SOM individual ancestries (alleles layer only) and three-layer SuperSOM species coefficients using the dataset generated in this study. Spearman’s and Pearson’s correlation coefficients are displayed. g) Genealogical divergence indices (gdi) calculated in HHSD. The dotted lines represent the suggested upper (split) and lower (merge) bounds for splitting populations into different species or merging them into a single species. Points represent the mean gdi values and lines represent the 95% highest probability density credible interval. “North” refers to the nuclear genomic clade containing all *P. regilla* and *P. sierra* samples. i) delimitR results for both the two-population and three-population analyses. In the model illustrations, dotted lines represent the halfway point between present day and the most recent divergence time. Arrows represent symmetrical gene flow between populations, and their position relative to the dotted line represents the timing of gene flow. Classification error and votes are displayed next to the models for both analyses. The models with the highest number of votes are indicated by bold text and red boxes, and their posterior probabilities (pp) are included. Alt text: SuperSOM BIC and change in BIC are displayed as box plots. Sampling frequency, layer weights, and variable importance plots are displayed as bar plots. In panel f, samples are represented by circles with sNMF admixture estimates on the x axis and DNA SOM individual ancestries and SuperSOM species coefficients on the y axes. HHSD mean gdi values are displayed as points on a plot. delimitR results are displayed in a table-like format. The left column contains a species tree for each tested model. The other columns contain classification error and votes for each model and delimitR analysis.

Details about prior settings and simulations are provided in Supplementary Materials. We simulated data under the nine models using 10,000 replicates each and built an RF classifier with the SFS bins as predictor variables, the model used to simulate data as the response variable, and 1000 decision trees. We used the RF_predict_abcrf function to apply the RF classifier to our dataset. The three-population models generated by delimitR do not account for both gene flow between *P. regilla* and *P. sierra* and divergence with gene flow between *P. hypochondriaca* and the “north” clade. We additionally ran delimitR with only two populations, “hypo” and “north”, because of the possibility of gene flow between *P. regilla* and *P. sierra* overpowering signals of gene flow between *P. hypochondriaca* and the “north” clade. We downprojected the SNP data to 142,260 (hypo,north) using easySFS (Gutenkunst et al. 2009; Overcast 2023). The population size prior, migration rate prior, and divergence time prior for *P. hypochondriaca* – “north” were kept the same. Divergence with gene flow and secondary contact were allowed, resulting in four possible models: one species (Model 1), two species without gene flow (Model 2), two species with secondary contact (Model 3), and two species with divergence with gene flow (Model 4) (Fig. 3h). We used 1000 decision trees when building the RF classifier.

To complement the other delimitation analyses, we used the program Hierarchical Heuristic Species Delimitation (HHSD) (Kornai et al. 2024). HHSD is a multispecies coalescent-based method that incorporates migration to calculate genealogical divergence indices (gdi), an estimated measure of genetic divergence between two populations resulting from gene flow and genetic isolation (Jackson et al. 2017; Leaché et al. 2019). HHSD consists of merge and split modes, in which taxa may be split or merged based on user-defined gdi thresholds. We assigned our samples to either “hypo”, “regilla”, or “sierra” using the same assignments as the delimitR analyses. Our input sequences consisted of the 549 loci from the ASTRAL dataset requiring at least 98% of individuals and no more than 5% gaps in the sequences. We set the merge threshold to <= 0.3 and the split threshold to >= 0.7, both of which are recommended thresholds for animals (Jackson et al. 2017; Kornai et al. 2024). We allowed migration to occur between “hypo” and “sierra” and between “regilla” and “sierra”. We performed initial testing with three different migration rate gamma priors to ensure robustness of the analyses. Because the results of these analyses did not substantially differ, we chose a migration rate prior of G(0.1, 20) to run the merge and split analyses for 500,000 generations each with a burn-in of 50,000 generations. We examined the trace plots and ESS values using Tracer v1.7.2 (Rambaut et al. 2018).

### Demographic Modeling

We used the program Genetic Algorithm for Demographic Model Analysis (GADMA) v2.0.3 to infer demographic history for the *P. regilla s.l.* complex (Noskova et al. 2020, 2023). Unlike delimitR which is limited to a set of user-defined models and symmetrical gene flow, GADMA uses a genetic algorithm to perform a global search of demographic parameters including the rate and direction of migration, divergence dating, and population size changes. We used easySFS to create an SFS downsampled to 10 samples per population (Gutenkunst et al. 2009; Overcast 2023), with samples assigned to the same “hypo”, “regilla”, and “sierra” populations as in the delimitR analyses. The moments engine was selected to infer demographic history (Jouganous et al. 2017), allowing for one time interval before and after each population split. Generation time was set to one year, and because germline mutation rate is not known for amphibians (Wang and Obbard 2023), we used the mean mutation rate for fish, 5.97e-9 (Bergeron et al. 2023), which is similar to rates used for reptiles and salamanders (Burbrink et al. 2022; Pyron et al. 2023a). Sequence length was estimated from the proportion of SNPs used to create the SFS multiplied by the total number of sites across all loci. We performed 20 repeats of this analysis and selected the top model based on the log likelihood. We later used Demes to plot the best model (Gower et al. 2022). Additionally, we reran this analysis using only two clades, *P. hypochondriaca* and “north”. Using easySFS, we downsampled the SFS to 20 samples per population. We set time intervals to allow one interval before the ancestral population diverged and three intervals after the divergence, with the goal of providing a more nuanced estimate for the timing of migration. All other parameters were kept the same as with the three-population analysis. To account for uncertainty in mutation rate and generation length parameters, we additionally calculated divergence times based on two additional mutation rates (Pyron et al. 2023a) and generation lengths that range from one year (Lemmon et al. 2007) up to five years (Mancini et al. 2025).

Lastly, we visualized gene flow across the landscape using Fast Estimation of Effective Migration Surfaces (FEEMS) (Marcus et al. 2021). FEEMS v1.0.0 estimates effective migration rates that deviate from expectations of IBD, allowing us to evaluate support for continuous geographic clines or discrete populations. If *P. regilla* – *P. sierra* or *P. hypochondriaca* – “north” splits are supported, we would expect to see reduced migration along the hypothesized geographic boundaries for these populations. Using the ingroup-only SNP dataset, we performed cross-validation runs for smoothing parameter (λ) values 0.01 – 20 and selected the λ value with the lowest cross-validation error as the “ideal” λ value. Larger values of the λ tuning parameter homogenize inferred migration weights, and lower values recover more fine-scale patterns with the potential for over-fitting. We fit the model using the ideal λ value (0.0495), as well as several other values of λ (0.815 and 10.0) to assess the impact of this parameter on results.

## Results

### Data Summary

Our sequenced dataset contained 215 *P. regilla s.l.* samples and three *P. cadaverina* samples because some samples were removed due to poor sequencing success. Samples had an average of 16 million reads per individual and 50× coverage per locus, and unaligned sequences were 199 bp in length. The original SNP dataset with one SNP per locus contained 11,643 SNPs, and the “ingroup-only” dataset contained 11,581 SNPs. The sequence dataset initially included 11,762 loci with an average alignment length of 225 (199 – 371) bp, prior to subsampling loci for ASTRAL and HHSD.

### Population Structure

Population structure inferred by sNMF and TESS3 showed similar patterns (Fig. 2a – f), with some incongruence when K = 3. For sNMF, the cross-entropy score stopped noticeably improving by K = 6 (Supplementary Fig. S1, S2); however, the automatic K-selection implemented in TESS3 selected K = 3 in most runs (Supplementary Fig. S3). For both methods, when K = 2, the samples clustered into a *P. hypochondriaca* group and a “north” group (Fig. 2a, c). We observed little admixture, primarily in localities on the range edges between *P. hypochondriaca* and the “north” group in southern California. When K = 3, the best sNMF run split the previous “north” group into an “interior” (Montana, eastern Washington) group and a “coastal” group, with extensive admixture between these two groups (Fig. 2b). TESS3 results for K = 3 were consistent with sNMF in about one-third of the runs (Fig. 2f). In another third of runs, TESS3 split the “north” group into a central California group and another group containing the remaining northern samples (Fig. 2e). The remaining third of runs split the “north” group into clusters aligning with the mitochondrial lineages for *P. regilla* and *P. sierra* (Fig. 2d). All three patterns showed extensive admixture between the subgroups within the “north” group.

The PCA formed two main clusters separated along the first axis of variance (Fig. 2g). One group included all samples associated with the geographic range for *P. hypochondriaca*. The second group contained “north” samples. New sampling localities occurring outside of the range boundaries inferred from Jadin et al. (2021) also fell within this group, including a sample from northern Tulare County, California between the known ranges for *P. hypochondriaca* and *P. sierra*. The separation between *P. regilla* and *P. sierra* samples on the second axis was not well-defined and rather displayed continuous variation between the two taxa.

### Phylogenetic Relationships

Phylogenies produced via SVDQuartets and ASTRAL supported different topologies that were discordant with mitochondrial lineages (Fig. 2h – l). In all trees, *P. hypochondriaca* and all “north” samples formed separate, well-supported clades. In the SVDQuartets tree, *P. regilla* was nested within *P. sierra* (Fig. 2h, Supplementary Fig. S4). In the ASTRAL trees, the 98% individuals and 5% missingness dataset supported clades for *P. regilla* and *P. sierra*, except with one *P. regilla* sample sister to all other samples within this “north” clade, and bootstrap support for *P. sierra* was weak (Fig. 2i, Supplementary Fig. S5). The datasets containing 98% individuals and 10% missingness and 92% individuals and 5% missingness supported *P. regilla* and *P. sierra* as separate clades (Fig. 2j – k, Supplementary Fig. S6, S7). The 92% individuals and 10% missingness dataset did not support distinct clades for *P. regilla* and *P. sierra* (Fig. 2l, Supplementary Fig. S8). The SplitsTree network analysis similarly revealed two distinct groups corresponding to *P. hypochondriaca* and “north” samples (Fig. 2m). Populations associated with *P. regilla* and *P. sierra* were not distinguishable within the “north” cluster.

### Species Delimitation

The three-layer SuperSOM using our nuclear dataset consistently sampled K = 2 across all 100 replicates (Fig. 3a). Cluster membership was delineated by the same *P. hypochondriaca* – “north” split observed in the population structure and phylogenetic analyses (Supplementary Fig. S9). SuperSOM species coefficients and DNA SOM ancestry coefficients were more sharply bimodal than individual ancestry coefficients estimated by SNPs alone in sNMF (Fig. 3f, Supplementary Fig. S10). The alleles layer had the highest layer weight and was much higher than the other two layers (Fig. 3b). Of the six bioclimatic variables, “Mean Temperature of Driest Quarter” had the highest variable importance, followed by “Mean Diurnal Range” (Fig. 3d). The four-layer SuperSOM using data from Vélez and Ingram (2025) also sampled K = 2 across all 100 replicates (Supplementary Fig. S11), though note that we did not have data from populations associated with *P. regilla*. Cluster membership delineated samples as expected (Supplementary Fig. S12), and species coefficients were completely bimodal (Supplementary Fig. S13), though this bimodal pattern could be influenced by the nature of mitochondrial inheritance. Alleles had the highest layer weight followed by traits, climate, and space respectively (Fig. 3c). The call variables in the traits layer with the highest variable importances were Pulse Rate and Number of Pulses in Phase 1 which were identified as the most different call properties between *P. hypochondriaca* and *P. sierra* by Vélez and Ingram (2025) (Fig. 3e). The climate variables with the highest variable importances, “Mean Temperature of Wettest Quarter” and “Precipitation Seasonality”, differed from the nuclear dataset SuperSOM (Fig. 3e).

The delimitR analysis with up to three populations supported Model 5, three species without gene flow, with 56.1% of the votes and a posterior probability of 0.61 (Fig. 3h). Model 9, three species with divergence with gene flow between *P. regilla* and *P. sierra*, received 43.0% of the votes, and most of the classification error for Model 5 was with Model 9 and vice versa. Models 1 – 4, representing only one or two species, received zero votes. Alternatively, the delimitR analysis allowing only up to two populations supported Model 2, two species without gene flow, with 47.90% of the votes and a posterior probability of 0.80 (Fig. 3h). Model 4, two species with divergence with gene flow, received 36.30% of the votes. Most of the classification errors for Model 2 and Model 4 were between these two models. The single species model, Model 1, received zero votes.

The HHSD merge analysis estimated gdi values of 0.51 (*P. regilla*) and 0.48 (*P. sierra*), putting these populations firmly in the “gray zone” of speciation and rejecting the merge (Fig. 3g). Estimated migration rates were asymmetrical, with a higher rate of migration (0.421 migrants per generation) from *P. sierra* into *P. regilla* and lower migration in the reverse (0.175 migrants per generation; Table 1).

**TABLE 1.**
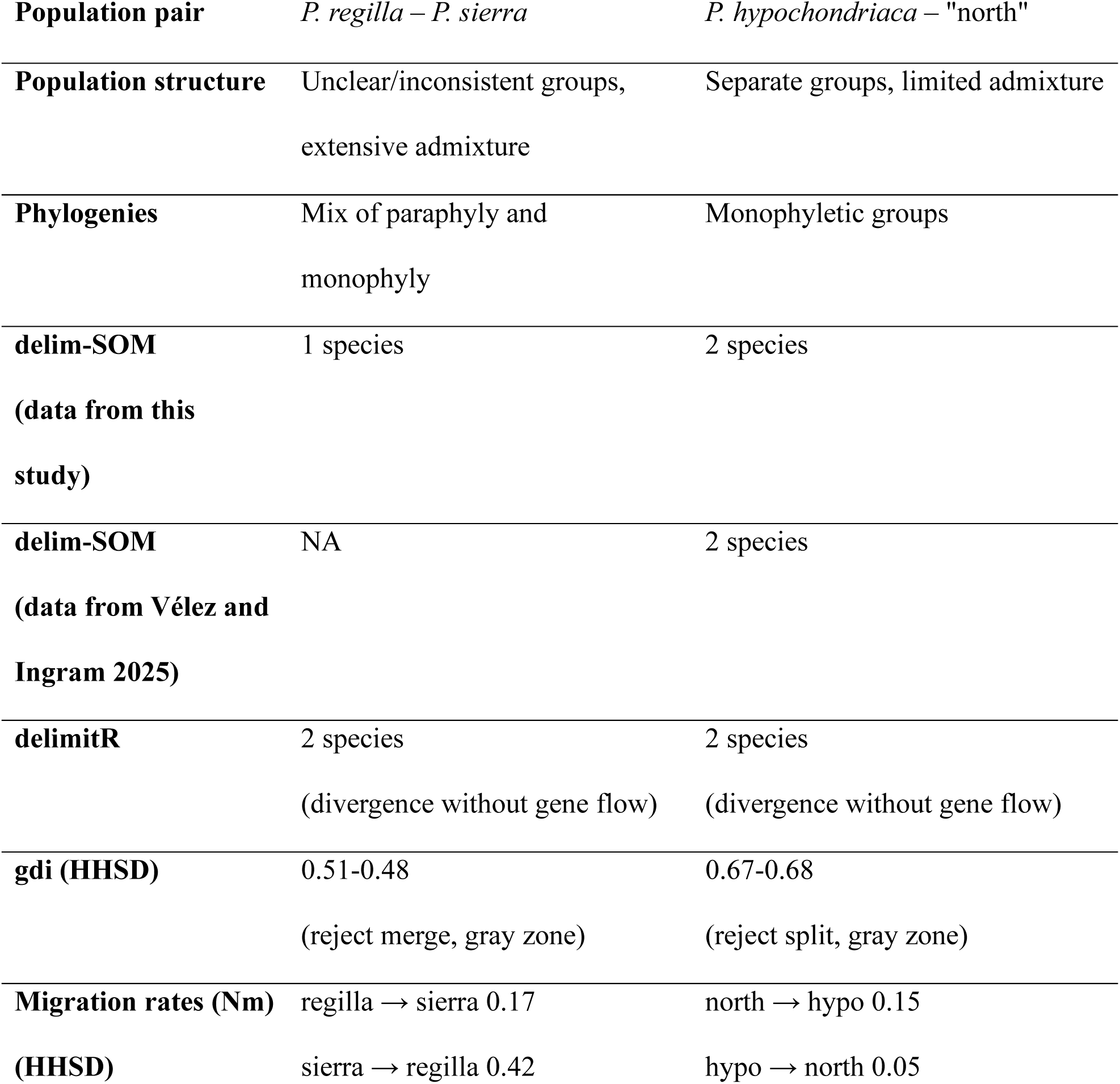
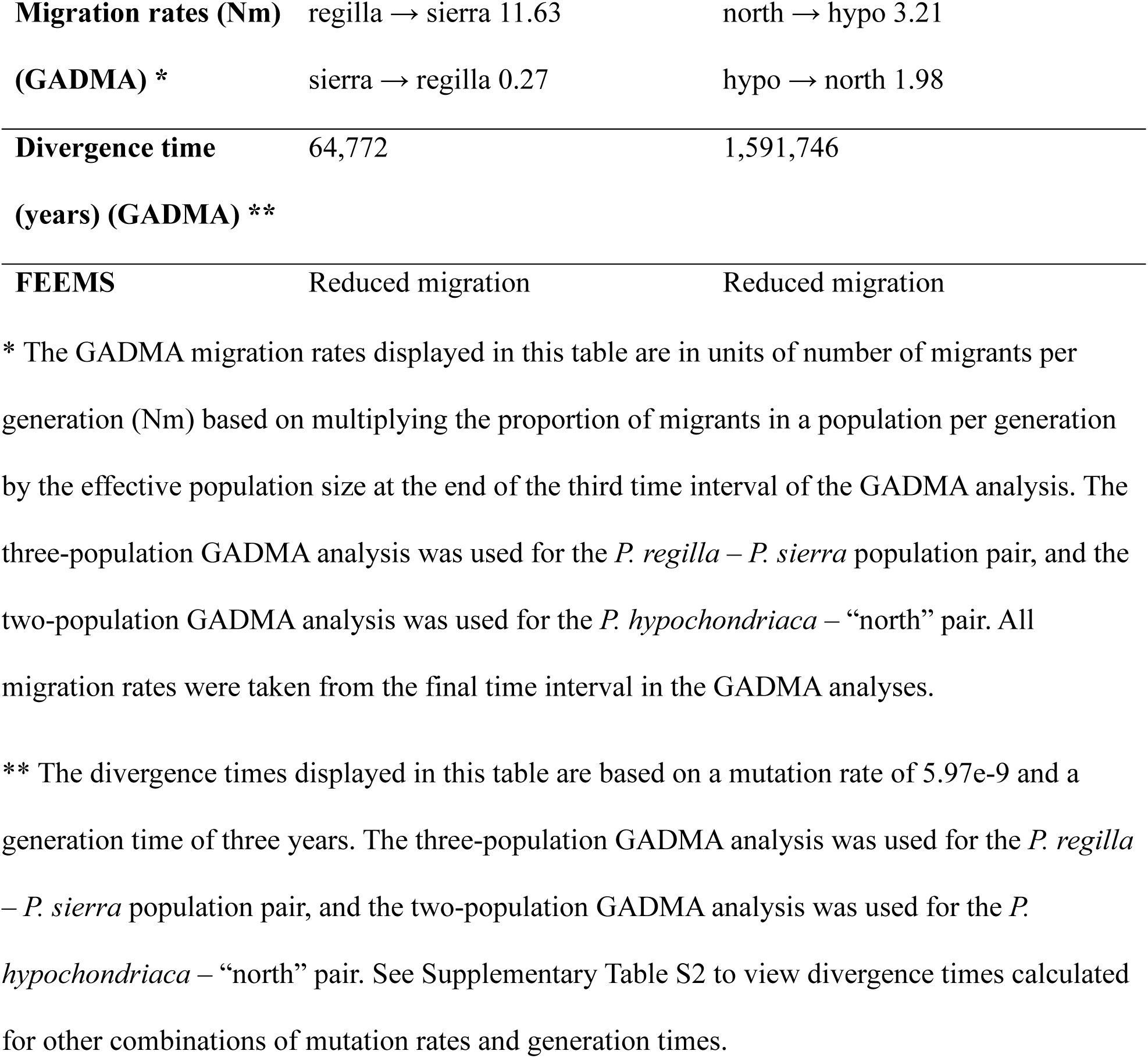
Summary of analyses results providing lines of evidence for species delimitation decisions for the *P. regilla* – *P. sierra* and *P. hypochondriaca* – “north” (*P. regilla* + *P. sierra*) population pairs.

The split analysis estimated gdi values of 0.67 (*P. hypochondriaca*) and 0.68 (“north”), rejecting the split (Fig. 3g); however, these values fall just below our specified guideline threshold (0.7) considered for species and the median gdi value for animals considered different species (0.68; Jackson et al. 2017), and the upper end of the 95% highest probability density credible interval for the “north” gdi value includes 0.7. Estimated migration rates between these two lineages were asymmetrical as well, with higher migration (0.152 migrants per generation) from “north” into *P. hypochondriaca* and lower migration in the reverse (0.045 migrants per generation; Table 1).

### Demographic Analyses

The best demographic model from GADMA for the three-population dataset inferred divergence between *P. hypochondriaca* and “north” populations 2,140,391 years ago and divergence between *P. regilla* and *P. sierra* 64,772 years ago, assuming a mutation rate of 5.97e-9 and a generation time of three years (Supplementary Table S2). Migration did not occur between *P. hypochondriaca* and the “north” group prior to the *P. regilla* – *P. sierra* divergence. After this divergence, migration is inferred between all three populations (Table 1, Supplementary Table S3). In contrast, the model inferred from the two-population dataset estimates the *P. hypochondriaca* – “north” divergence occurring 1,583,788 years ago, under the same mutation rate and generation length assumptions. Migration was not inferred until 46,953 years ago, starting with unidirectional migration from “north” into *P. hypochondriaca*, and then bidirectional migration beginning 26,376 years ago (Table 1, Supplementary Table S3).

Our FEEMS analyses revealed similar overall patterns across the different values of λ, though lower values of λ were potentially overfit, inferring finer scale patterns that were not informative for our purposes (Fig. 4, Supplementary Fig. S14). Effective migration rates were lowest along the *P. hypochondriaca* – *P. sierra* boundary (Fig. 4). Migration rates were also reduced near the *P. regilla* – *P. sierra* boundary and between “curta” populations o *P. hypochondriaca* in Baja California Sur and the remaining *P. hypochondriaca* populations.

**FIGURE 4.**
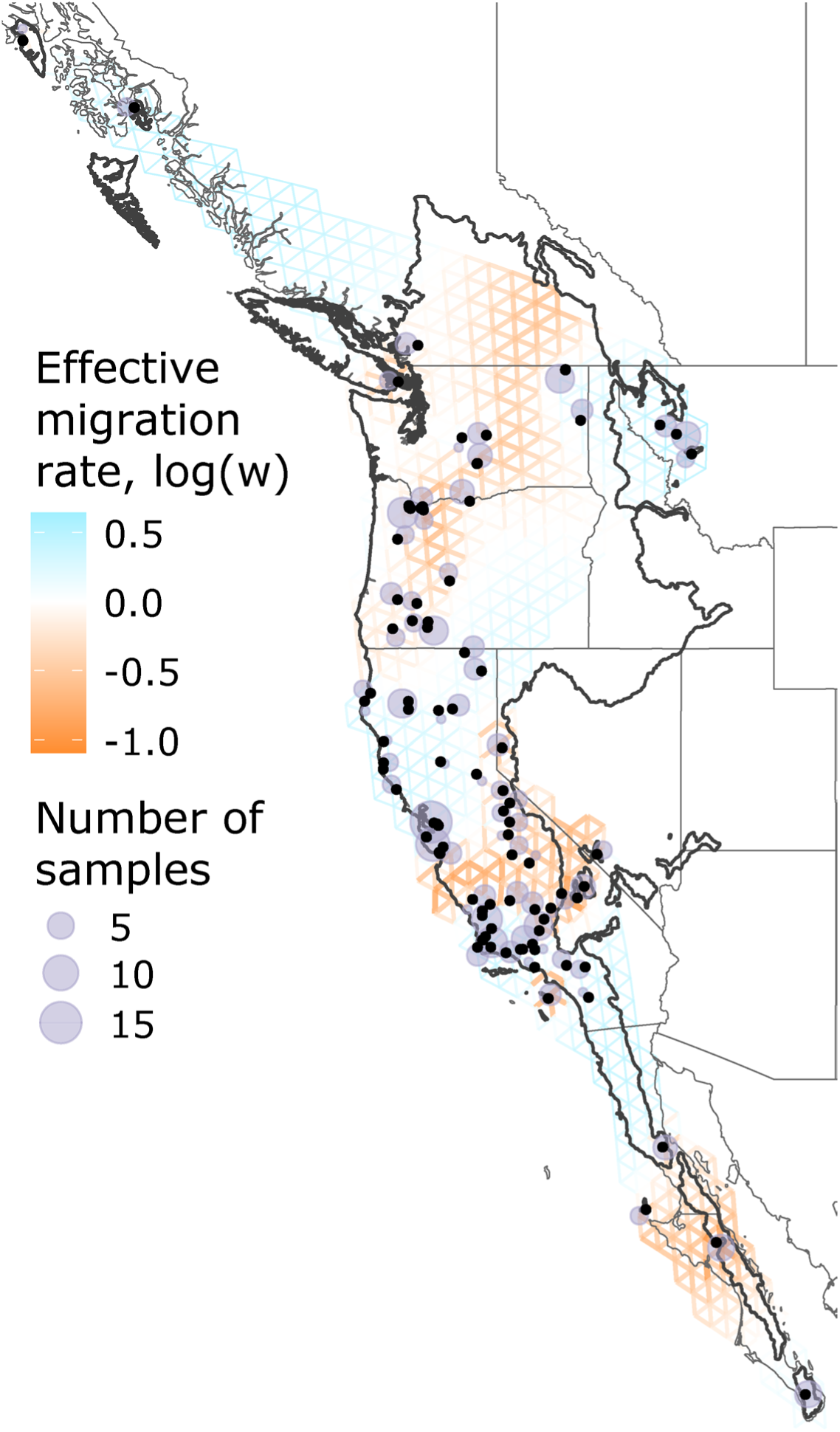
FEEMS results when the smoothing parameter (λ) = 10.0. Line thickness and color represent the effective migration rate, with orange and thicker lines representing lower effective migration rates, and blue and thinner lines representing higher effective migration rates. The number of samples for each node is represented by the size of the purple circle. Points of the original samples are added as black dots. The range of the *P. regilla sensu lato* complex is outlined in black. Alt text: The FEEMS map shows a grid of demes across the landscape, representing effective migration surfaces.

## Discussion

In this study, we aimed to resolve inconsistencies in the systematic treatment of *Pseudacris regilla s.l.*, an iconic frog species complex with variable morphology and broad geographic and environmental distributions. Using thousands of nuclear loci, we discovered genetic population clusters, inferred phylogenetic relationships, implemented species delimitation approaches, and elucidated the demographic history of divergence and gene flow between lineages. We found consistent support for *P. hypochondriaca* and “north” lineages, but support for *P. regilla* and *P. sierra* as separate species was inconsistent. We discuss species delimitation in the gray zone of speciation, particularly the use of multiple analytical methods to reveal incongruence in results, with the *P. regilla s.l.* complex as a case study. We additionally make recommendations for taxonomic treatment and identification in this system and suggestions for further directions of study.

### Species Delimitation in the Gray Zone

Species delimitation remains challenging even with expansive geographic and genomic sampling, particularly for lineages that fall within the “gray zone” of speciation. A typical species delimitation approach is to first discover putative groups, often through *a priori* hypotheses and population structure analysis, and then perform group validation using species delimitation methods (Chambers et al. 2025b; Pavón-Vázquez et al. 2026). However, models are simplified versions of complex biological processes, and incongruence between methods may result from assumption violations or differences in statistical power to detect independent lineages (Carstens et al. 2013; Jacobs et al. 2018). Incongruence may also be attributed to differences in signal and history between data types, exemplified by patterns such as mitonuclear discordance (Toews and Brelsford 2012). Congruence in species delimitation results across multiple methods provides stronger confidence in species delimitation decisions, whereas incongruence requires closer scrutiny about its potential causes and should lead to more conservative inferences (Carstens et al. 2013; Kuchta et al. 2016). We took this approach of using multiple methods and different parameters in testing our species hypotheses, allowing us to detect inconsistencies across datasets and methods and evaluate uncertainty in our data and prior knowledge of the study system.

A common source of incongruence is found between gene trees and species trees, often due to processes such as incomplete lineage sorting (ILS) or introgression (Toews and Brelsford 2012; Steenwyk et al. 2023). We observed this discordance with *P. regilla s.l.*, where mitochondrial phylogenies inferred a different topology than the species trees inferred by nuclear loci in our study and previous studies (Recuero et al. 2006; Barrow et al. 2014; Banker et al. 2020; Jadin et al. 2021; Vélez and Ingram 2025; see note about nuclear monophyly in Supplementary Materials). With multiple nuclear loci, assumptions behind species tree methods and how they handle these patterns can also cause incongruence between species tree methods. The popular species tree method ASTRAL (Zhang et al. 2018) can account for ILS, but it is reliant on the accuracy of the input gene trees and is prone to error with gene trees inferred from shorter sequences (Chou et al. 2015). Alternatively, the usage of SNPs by SVDQuartets (Chifman and Kubatko 2014, 2015) overcomes the limitation of short sequences lengths but can be misled by high levels of ILS (Chou et al. 2015). Incomplete lineage sorting can result from recent divergence times (Maddison and Knowles 2006), which could explain the discrepancies observed between methods for the *P. regilla* and *P. sierra* relationship. Neither of these methods account for introgression, so utilizing methods that infer non-treelike patterns, such as SplitsTree networks (Huson and Bryant 2024), can help clarify these relationships.

Incongruent results among species delimitation methods may be explained by different model assumptions and statistical power to detect independent lineages (Carstens et al. 2013; Jacobs et al. 2018). Our use of delim-SOM (Pyron et al. 2023b) allowed us to integrate multiple data types and both discover and validate groups without making biological assumptions about the study system. This method invariably delimited two species. However, model-based species delimitation approaches are also important to put the process of divergence in a biologically meaningful context. We compared models that incorporate divergence and gene flow using delimitR (Smith and Carstens 2020) and recovered the best models as ones that assume no gene flow between populations and support up to three species (Fig. 3h). However, we encountered high classification errors in our delimitR analyses which could be attributed to recent divergence times or misspecification of priors (Smith and Carstens 2020). A third species delimitation approach is to visualize where in the “gray zone” of speciation populations may fall, which we implemented by calculating gdi values in HHSD (Jackson et al. 2017; Leaché et al. 2019; Kornai et al. 2024). Gray zone populations will likely fall between the upper and lower guidelines for consideration of separate or single species, but values closer to these thresholds provide stronger support for or against separate species on this spectrum (Jackson et al. 2017). Our *P. regilla s.l.* gdi values placed *P. regilla* and *P. sierra* centrally in the gray zone, whereas *P. hypochondriaca* and “north” were placed at or just below the median gdi for animal species (Fig. 3g, Table 1). The *P. regilla s.l.* populations are considered gray zone taxa by this measure, but this method demonstrated these populations fall within the gray zone of speciation to different degrees, with *P. hypochondriaca* and “north” closer to species.

### Mechanisms of Divergence

Understanding the mechanisms underlying the divergence process can provide important context and biological rationale for species delimitation decisions. Geological processes, climatic variation, and reproductive behavior are likely mechanisms contributing to the *P. hypochondriaca* – “north” divergence and secondary contact. Intermediate values of mutation rate and generation length suggest *P. hypochondriaca* and “north” diverged around 1.6 Ma. This date is younger than the *Pseudacris regilla s.l.* – *P. cadaverina* divergence time estimate (3.3 Ma) but older than the *P. hypochondriaca* – *P. sierra* estimate (0.8 Ma) using mitochondrial data (Jadin et al. 2021), but caution should be used when interpreting divergence dates when mutation rates and generation length are unknown (see Supplementary Table S2 for alternative estimates). The southern Sierra Nevada mountains, the inferred boundary between *P. hypochondriaca* and “north” east of the Central Valley, experienced rapid uplift during the late Pliocene or Quaternary (Figueroa and Knott 2010) during the inferred *P. hypochondriaca* – “north” divergence time. Our data show the boundary between *P. hypochondriaca* and “north” could also be associated with the Monterey Bay on the west side of the Central Valley, which is a reported phylogeographical break in other herpetofauna (Calsbeek et al. 2003; Rissler et al. 2006). The Central Valley was an ancient embayment with drainage into Monterey Bay until 2 Ma and a subsequent emergent basin until 0.6 Ma (Sarna-Wojcicki et al. 1985). Despite the embayment preceding the *P. hypochondriaca* – “north” divergence, it may have reduced gene flow between these lineages and maintained their divergence.

In conjunction with geological changes in the Sierra Nevada, climatic changes may have impacted these lineages differently. Climatic niche models by Jadin et al. (2021) showed some niche overlap and differentiation between *P. hypochondriaca* and *P. sierra* during the Last Interglacial (LIG), Last Glacial Maximum (LGM), and present with ranges that were relatively stable through time. Notably, the San Joaquin Valley appears unfavorable to *P. sierra* in the LIG and present niche models, and the Sierra Nevada appears unfavorable to *P. hypochondriaca* in all three time periods. While these time periods are younger than the *P. hypochondriaca* – “north” divergence, these niche models suggest potential climatic niche differences that could have developed earlier in the past. The Sierra Nevada mountains underwent glaciations throughout the Pleistocene (Bateman 1968) which could have isolated these lineages in separate refugia and allowed them to diverge in allopatry, with recent secondary contact after glaciation. Niche overlap between *P. hypochondriaca* and *P. sierra* in the San Joaquin Valley during the LGM could also explain the recent secondary contact inferred between *P. hypochondriaca* and “north” starting approximately 26,000–47,000 years ago (Supplementary Table S2–S3).

Reproductive call differentiation may also be an important contributing mechanism of divergence between *P. hypochondriaca* and “north”. In frogs, reproductive call differences are recognized as an important mechanism that can promote or reinforce reproductive isolation during the speciation process (Blair 1964; Hoskin et al. 2005; Wilkins et al. 2013). Mating calls from *P. hypochondriaca* have significantly lower pulse rate and fewer pulses than “north” populations in California (Vélez and Ingram 2025). The exact reason and historical timing of this differentiation is unknown, but the authors hypothesized that this change in call properties could have occurred if *P. hypochondriaca* shared Pleistocene refugia with *P. cadaverina*, driving the differentiation in acoustic signaling through reproductive character displacement (Vélez and Ingram 2025). The authors also noted that these acoustic properties exhibited significant phylogenetic signal in their data, implying that these call traits evolved in step with the initial lineage divergence. These acoustic properties are known to affect species recognition in other *Pseudacris* species (Lemmon 2009), though the extent to which females of either *P. hypochondriaca* or “north” discriminate between these calls is unknown. Future studies examining female mate choice between lineages and historical niche modeling for *P. hypochondriaca* and *P. cadaverina* could reduce these knowledge gaps.

For the *P. regilla* and *P. sierra* divergence, geological and climatic explanations have been investigated previously. The Klamath mountains have been noted as occurring along the southwestern edge of the *P. regilla* range (Jadin et al. 2021), and while they might reduce gene flow, their age significantly predates our estimated *P. regilla* – *P. sierra* divergence (Harper and Wright 1984), making these mountains a possible barrier to gene flow, but an unlikely contributor to the initial divergence. Before the Pleistocene, the rain shadow from the major uplift of the Cascade Mountains led to the xerification of the Columbia Plateau (Graham 1999; Brunsfeld et al. 2001). A pattern of disjunct Pacific coast and interior Rocky Mountain populations on either side of the Columbia Plateau is documented in other amphibians (Brunsfeld et al. 2001) with evidence of ancient vicariance (Carstens et al. 2005), and the Columbia Plateau has been proposed as a potential barrier to dispersal between *P. regilla* and *P. sierra* based on mitochondrial divergence time estimates (∼1.4 – 3.3 Ma; Ripplinger and Wagner 2004; Jadin et al. 2021). However, our nuclear data support a much younger divergence time; under intermediate divergence time assumptions, divergence between *P. regilla* and *P. sierra* began approximately 65,000 years ago during the Last Glacial Period. Niche models hindcasted onto the LIG (∼120,000–140,000 years ago, before the nuclear divergence time between *P. regilla* and *P. sierra*) and LGM (∼22,000 years ago, after their divergence) infer that *P. sierra* had a relatively stable range with only recent expansion into the northwest portion of its range, whereas the *P. regilla* range contracted during the LGM, and the overlap between niches appears limited (Jadin et al. 2021). This range contraction between the LIG and the LGM could have resulted in separate refugia for *P. regilla* and *P. sierra*, allowing time for divergence in isolation. Migration estimates indicated proportionally low but extremely asymmetrical rates of gene flow, but because we did not infer more precise timing of gene flow for this population pair, it is unclear if subsequent warming with potential range expansion led to secondary contact and gene flow, or if other scenarios such as divergence with gene flow or pulses of gene flow were occurring. Acoustic differentiation that may have led to divergence between *P. regilla* and *P. sierra* is currently unknown. Populations of *P. regilla* from San Juan Island in Washington reportedly have similar pulse rates to California populations of *P. sierra* (Rose and Brenowitz 2002; Vélez and Ingram 2025); however, male acoustic signaling and female preferences may differ between allopatric and sympatric populations (Lemmon 2009), so further study is needed to characterize acoustic signaling for *P. regilla* and *P. sierra* throughout their ranges and at contact zones.

### Taxonomic Recommendations and Species Identification

The *P. hypochondriaca* – “north” pair displayed a consistent pattern of two distinct groups across methods, and the gdi values, the main source of incongruence, still found them to fall at or just below the median value for species (Table 1). Demographic models suggest these two lineages diverged without gene flow about 1.6 Ma with recent, but limited, secondary contact within the past few tens of thousands of years (Table 1, Supplementary Table S2–S3). Niche models suggest these two lineages inhabit different climatic niches (Jadin et al. 2021), and acoustic differentiation also supports two separate groups and may act as a potentially important reproductive isolating mechanism (Vélez and Ingram 2025). We recommend retaining the recognition of Baja California Treefrogs (*Pseudacris hypochondriaca*) as a distinct species from the “north” lineage that contains *P. regilla* and *P. sierra*. Based on our current sampling, *Pseudacris hypochondriaca* inhabits southern California, southern Nevada, and the Baja California peninsula in Mexico (Fig. 1d).

In contrast, *P. regilla* and *P. sierra* displayed incongruent results across methods. Detection of separate groups, phylogenetic relationships, and the inference of separate species in species delimitation analyses were inconsistent (Table 1). This pair fell within the middle of the “gray zone” for the HHSD analysis. Our models suggest that *P. regilla* and *P. sierra* experienced recent divergence about 65,000 years ago with unknown timing of gene flow. We recommend that *P. regilla* and *P. sierra* be considered one species. *Pseudacris sierra* should be synonymized with *P. regilla*, with *P. regilla* taking priority (Baird and Girard 1852); hereafter, *P. regilla* refers to the “north” nuclear genomic lineage, containing all samples previously assigned to *P. regilla* and *P. sierra*. Thus, Pacific Treefrogs (*Pseudacris regilla*) occur throughout northern California, parts of central California, northern Nevada, Idaho, Montana, Oregon, Washington, Alaska, and British Columbia in Canada (Fig. 1d). Future investigations of contact zone dynamics and acoustic properties may provide lines of evidence to support species status. For example, asymmetrical gene flow or hybridization, which we inferred between *P. hypochondriaca* and *P. regilla* but more drastically between the two *P. regilla* populations, can be indicative of reinforcement via reproductive character displacement in female preference (Lemmon and Juenger 2017). However, current evidence examined in this study indicates the previously described mitochondrial lineages within *P. regilla* do not align with consistent species-level nuclear genomic variation.

Identification between *P. hypochondriaca* and *P. regilla* can be achieved in several ways. The best identifiers in the field are geographic location and advertisement calls. At present, we do not have evidence of sympatry at sampling locations, thus individuals found within the known sampling boundaries most likely correspond to the appropriate species based on location (Fig. 1d). However, a sampling gap remains between these species, particularly in the Central Valley, and human-mediated translocation of individuals is possible. Examining acoustic properties of advertisement calls such as the pulse rate or number of pulses can serve as another relatively reliable identification method. Note that temperature must be considered during identification because it influences these acoustic traits (Gayou 1984; Vélez and Ingram 2025), and that *P. regilla* acoustic data was only assessed in California for populations previously attributed to *P. sierra* (Vélez and Ingram 2025). In a lab environment, tissue samples collected from these frogs can be sequenced for cytb for identification. To our current knowledge, there is no evidence of mitochondrial introgression between *P. hypochondriaca* and *P. regilla*, but fine-scale sampling at contact zones is needed to confirm this assumption. Ideally, morphological characteristics would make identification straight-forward. Jameson et al. (1966) described morphological variation across this species complex; however, the taxonomic assignments in their study do not align with our current understanding of the geographic distribution of these species (Fig. 1a). Georeferenced museum specimens represent an untapped opportunity for investigating whether there are morphological differences between *P. hypochondriaca* and *P. regilla*.

### Concluding Remarks

Ultimately, species delimitation in the gray zone requires making subjective decisions informed by the extent of our current knowledge. Congruence across methods and datasets provides stronger support for separate species, whereas incongruence requires careful consideration. Utilizing multiple methods for species delimitation is thus essential to detect potential incongruence and inform decisions. Importantly, understanding the divergence mechanisms operating within the study system can help provide biological context to the speciation process and additional lines of evidence for species delimitation decisions.

Our genomic analyses indicate that the iconic “Hollywood frog” is comprised of not one or three species, but two species that diverged between the Early to Middle Pleistocene with recent secondary contact. We proposed several hypotheses about the mechanisms of divergence that remain to be investigated with additional sampling, analyses, and behavioral experiments, particularly to understand the exact location and nature of the contact zones between *P. hypochondriaca* and *P. regilla* and between the two *P. regilla* lineages. Dynamics of the *P. hypochondriaca* – *P. regilla* secondary contact zone could impact the evolutionary trajectories of these lineages, with potential for lineage fusion or reinforcement (Abbott et al. 2013; Lemmon and Juenger 2017). Future work with fine-scale sampling will help determine more precise geographic range boundaries between species, investigate potential gene flow between these species at a local scale, and further elucidate the history and mechanisms of isolation.

## Supporting information

Supplementary Methods, Notes, References, and Figures

## Acknowledgements

We thank the University of New Mexico Center for Advanced Research Computing, supported in part by the National Science Foundation, for providing the high-performance computing and large-scale storage resources used in this work. We thank Curtis Schmidt and Joseph T. Collins (FHSM), Carol Spencer and Jimmy McGuire (MVZ), Sharon Birks and Adam Leaché (UWBM), and Andrés Lopez and Kyndall Hildebrandt (UAM) for tissue loans. We thank Moses Michelsohn, Luis Amador, and Dani Wiley for comments that helped improve this manuscript. We also thank the two anonymous reviewers, Associate Editor Danielle Edwards, and Editor-in-Chief Robert C. Thomson for helpful comments that greatly improved this manuscript.

## Data Availability

All supplementary materials can be accessed through the online version of this article. Original data and scripts necessary to reproduce the analyses reported in this study can be accessed through the Dryad link: https://doi.org/10.5061/dryad.1jwstqk94.

