## Supplementary Methods, Notes, References, and Figures for "The Tangled History and Taxonomy of an Iconic Chorus Frog Complex Clarified using Genomic Analyses"

Finally, we ran a Principal Component Analysis (PCA) to visualize population clusters along the two axes explaining the highest proportion of variance in the data. We used PGDSpider v3.0.0.0 to convert the VCF file to STRUCTURE format (Lischer and Excoffier 2012). Settings were left at defaults except for changing the datatype to “SNP”. In R, we used the read.structure function in the adegenet package (Jombart 2008; Jombart and Ahmed 2011) to read in the structure file and then converted the genind object to a genlight object using the gi2gl function in the dartR package (Gruber et al. 2018; Mijangos et al. 2022). We used the gl.pcoa and gl.pcoa.plot functions (Gruber et al. 2018; Mijangos et al. 2022) to run the PCA and visualize the results in a biplot.

Our first SuperSOM used our nuclear genomic dataset generated in this study and included alleles, space (latitude, longitude, elevation), and climate (bioclimatic variables) as input layers. We read in our structure file of SNPs using the read.structure function and then calculated allele frequencies using the makefreq function, both in the adegenet package (Jombart 2008; Jombart and Ahmed 2011). We retrieved elevation data from coordinates using the get_elev_point function in the elevatr package (Hollister et al. 2025). We downloaded bioclimatic data from the WorldClim version 2.1 dataset at 2.5 arcminutes (Fick and Hijmans 2017) and obtained the values for each specimen coordinates using the extract function in the terra package (Hijmans et al. 2026). We retained only six bioclimatic variables after identifying correlated variables using the corSelect function in the fuzzySim package (Barbosa 2015). We normalized the space and climate data to a scale of 0–1 using the delim-SOM minmax function (Pyron et al. 2023b) and the dplyr mutate function (Wickham et al. 2026) to put them on the same scale as the allele frequency data. We created an output grid using the somgrid function in the delim-SOM package (Pyron et al. 2023b) and then ran a three-layer SuperSOM using the Climate.SOM function (Pyron et al. 2023b) including 100 replicates and 100 steps. We also ran a DNA SOM using only the alleles layer in the DNA.SOM function (Pyron et al. 2023b) to make direct comparisons to our sNMF analysis.

Our second SuperSOM used data from Vélez and Ingram (2025) including alleles (mitochondrial cytochrome *b* sequences), space (latitude, longitude, elevation), climate (bioclimatic variables), and traits (reproductive call properties). We downloaded 31 of the mitochondrial sequences from NCBI GenBank (Clark et al. 2016) and aligned them in AliView (Larsson 2014) using Muscle version 3.8.425 (Edgar 2004). We imported the alignment into R using the read.alignment function in the seqinr package (Charif and Lobry 2007), converted the alignment to a genind object using the alignment2genind function in the adegenet package (Jombart 2008; Jombart and Ahmed 2011), and calculated allele frequencies using the makefreq function in the adegenet package (Jombart 2008; Jombart and Ahmed 2011). We tried coding each haplotype as its own variable in the alleles layer, but there were too many unique haplotypes to produce signal in the results. We therefore treated variations in the sequence alignment as SNPs but acknowledge the potential of introducing pseudo-replication. As input for the traits layer, we chose the mean for each of nine call variables from the locality where each sequence was collected in Vélez and Ingram (2025). We additionally used the latitude, longitude, elevation, and bioclimatic variables provided in Vélez and Ingram (2025). We retained the same six bioclimatic variables as in the first SuperSOM. We normalized all space, climate, and trait variables to a scale of 0–1 as described previously. We ran a four-layer SuperSOM using the Trait.SOM function (Pyron et al. 2023b) with the same input grid, number of replicates, and number of steps as in our first SuperSOM.

To incorporate demographic scenarios involved in lineage formation, we used the species delimitation R package delimitR (Smith and Carstens 2020). delimitR uses fastsimcoal2 simulations to model different combinations of number of species, presence and timing of gene flow, and number of migration edges (Excoffier et al. 2013; Smith and Carstens 2020). The program then uses a random forest (RF) classifier to select which model best reflects the data. We assigned our samples to either “hypo”, “regilla”, or “sierra” based on their positions in the ASTRAL trees, though we note that the evidence for three groups, especially the designations for *P. regilla* and *P. sierra*, are equivocal considering the inconsistent results for K = 3 in the population structure analyses, the differing tree topologies, and the lack of detection for K = 3 by delim-SOM. These assignments were kept to evaluate the previous species hypotheses generated by Recuero et al. (2006) and Jadin et al. (2021) and the demographic history of this group using a model-based method (assignments indicated in Supplementary Table S1). The ingroup-only VCF file was converted to the multi-dimensional site frequency spectrum (mSFS) using easySFS v0.0.1 and downprojected to 76,188,142 (regilla,sierra,hypo) because these values maximized the number of segregating sites (Gutenkunst et al. 2009; Overcast 2023). We changed the first cell in the output file to “0” to remove monomorphic sites. In delimitR, we used the setup_fsc2 function to create nine models (Fig. 3i): one species (Model 1), two species without gene flow (*P. hypochondriaca* and “north”; hereafter, “north” refers to the clade containing all *P. regilla* and *P. sierra* samples) (Model 2), two species with secondary contact (Model 3), two species with divergence with gene flow (Model 4), three species without gene flow (Model 5), three species with secondary contact between *P. regilla* and *P. sierra* (Model 6), three species with secondary contact between *P. hypochondriaca* and *P. sierra* (Model 7), three species with secondary between *P. regilla* and *P. sierra* and between *P. hypochondriaca* and *P. sierra* (Model 8), and three species with divergence with gene flow between *P. regilla* and *P. sierra* (Model 9). Because information is lacking on population sizes for the *P. regilla s.l.* complex, we used a broad uniform population size prior of 500 to 1,000,000 for all populations. Divergence time priors were determined using estimates from Jadin et al. (2021). For the *P. hypochondriaca* – “north” split, we used the upper confidence interval for the *P. cadaverina* – *P. regilla s.l.* complex split, approximately 6,000,000 years, as the upper threshold for this prior, and the lower estimate for the *P. hypochondriaca* – *P. sierra* split, 53,000 years, for the lower threshold. Because Jadin et al. (2021) considered *P. hypochondriaca* and *P. sierra* to be sister species rather than *P. regilla* and *P. sierra*, we did not have an estimate for the *P. regilla* – *P. sierra* split and instead used the upper estimate for the *P. regilla s.l.* complex, 2,624,000 years, as the upper threshold for this prior, and the lower threshold was set to 500 to represent the very recent past. The divergence time prior uses generation time rather than years; however, generation time has not been calculated for the *P. regilla s.l.* complex, so we assumed a probable range of 1–3 years for the generation time (Lemmon et al. 2007; Ethier et al. 2021), resulting in a final uniform prior of 15,000 to 6,000,000 for the *P. hypochondriaca* – “north” split and 500 to 3,000,000 for the *P. regilla* – *P. sierra* split. Because the divergence time priors overlap, an extra setting was defined to make sure the *P. hypochondriaca* – “north” split divergence time was oldest. We used a broad uniform migration rate prior of 0.000005 to 0.05. We used the fastsimcoalsims function to simulate data under the nine models using 10,000 replicates each. We summarized the mSFS using five classes with the makeprior function and removed rows with zero variance using the Prior_reduced function. The RF_build_abcrf function was used to build an RF classifier with the bins of the binned SFS as the predictor variables, the model used to simulate data as the response variable, and 1000 decision trees. We lastly used the prepobserved function to convert our observed data into the correct format and then used RF_predict_abcrf to apply the RF classifier to our dataset.

The three-population models generated by delimitR do not account for both gene flow between *P. regilla* and *P. sierra* and divergence with gene flow between *P. hypochondriaca* and the “north” clade. We additionally ran delimitR with only two populations, “hypo” and “north”, because of the possibility of gene flow between *P. regilla* and *P. sierra* overpowering signals of gene flow between *P. hypochondriaca* and the “north” clade. We downprojected the SNP data to 142,260 (hypo,north) using easySFS (Gutenkunst et al. 2009; Overcast 2023). The population size prior, migration rate prior, and divergence time prior for *P. hypochondriaca* – “north” were kept the same. Divergence with gene flow and secondary contact were allowed, resulting in four possible models: one species (Model 1), two species without gene flow (Model 2), two species with secondary contact (Model 3), and two species with divergence with gene flow (Model 4) (Fig. 3j). We used 1000 decision trees when building the RF classifier.

To complement the other delimitation analyses, we used the program Hierarchical Heuristic Species Delimitation (HHSD) (Kornai et al. 2024). HHSD is a multispecies coalescent-based method that incorporates migration to calculate genealogical divergence indices (gdi), an estimated measure of genetic divergence between two populations resulting from gene flow and genetic isolation (Jackson et al. 2017; Leaché et al. 2019). HHSD consists of merge and split modes, in which taxa may be split or merged based on user-defined gdi thresholds. We assigned our samples to either “hypo”, “regilla”, or “sierra” using the same assignments as the delimitR analyses. Our input sequences consisted of the 549 loci from the ASTRAL dataset requiring at least 98% of individuals and no more than 5% gaps in the sequences. We set the merge threshold to <= 0.3 and the split threshold to >= 0.7, both of which are recommended thresholds for animals (Jackson et al. 2017; Kornai et al. 2024). We allowed migration to occur between “hypo” and “sierra” and between “regilla” and “sierra”. We performed initial testing with three different migration rate gamma priors to ensure robustness of the analyses: G(0.1, 20) with a mean of 0.005, G(0.1, 10) with a mean of 0.01, and G(2, 20) with a mean of 0.1. Each of the split and merge analyses ran for 200,000 generations with a burn-in of 20,000 generations. Because the results of these analyses did not substantially differ, we chose a migration rate prior of G(0.1, 20) to run the merge and split analyses for 500,000 generations each with a burn-in of 50,000 generations. We examined the trace plots and ESS values using Tracer v1.7.2 (Rambaut et al. 2018).

Lastly, we visualized gene flow across the landscape using Fast Estimation of Effective Migration Surfaces (FEEMS) (Marcus et al. 2021). FEEMS v1.0.0 estimates effective migration rates that deviate from expectations of IBD, allowing us to evaluate support for continuous geographic clines or discrete populations. If *P. regilla* – *P. sierra* or *P. hypochondriaca* – “north” splits are supported, we would expect to see reduced migration along the hypothesized geographic boundaries for these populations. Following Chambers et al. (2025b), we used BCFTools v1.21 (Danecek et al. 2021) to filter out rare variants and exclude SNPs in which over 50% of samples had a missing genotype from our ingroup-only VCF file. Plink v1.9.0-b.7.7 (Purcell et al. 2007) was then used to convert the data into input BIM, BED, and FAM files. We used the dggridR package (Barnes and Sahr 2024) to create a discrete grid. With FEEMS, we performed cross-validation runs for smoothing parameter (λ) values 0.01 – 20 and selected the λ value with the lowest cross-validation error as the “ideal” λ value. Larger values of the λ tuning parameter homogenize inferred migration weights, and lower values recover more fine-scale patterns with the potential for over-fitting. We fit the model using the ideal λ value (0.0495), as well as several other values of λ (0.815 and 10.0) to assess the impact of this parameter on results.

Notes

Banker et al. (2020) and Barrow et al. (2014) inferred the three taxa in the *P. regilla s.l.* complex to be non-monophyletic based on nuclear DNA. However, one *P. hypochondriaca* sample (MHP14886) was erroneously labeled as *P. sierra* because of an error in Fig. 1 from Recuero et al. (2006), where samples from populations 17 and 18 were incorrectly grouped with *P. sierra* on the sampling map but correctly labeled in the phylogeny. If corrected, all three taxa are monophyletic in Banker et al. (2020) and sometimes monophyletic in Barrow et al. (2014) depending on the dataset and analysis. In Jadin et al. (2021), Recuero et al. (2006) samples from localities 17 and 18 were also mislabeled as *P. sierra* on the map in Fig. 1 and in the table Appendix S2. One *P. sierra* sample (GenBank #KJ536203) used in Jadin et al. (2021) was erroneously attributed to coordinates corresponding to San Luis Obispo County. This sample is originally from Mendocino County (ECM2694; Barrow et al. 2014). When correcting the identifications and localities of these samples, this changes the known mitochondrial lineage in San Luis Obispo County from *P. sierra* into *P. hypochondriaca*.

Supplementary Figures
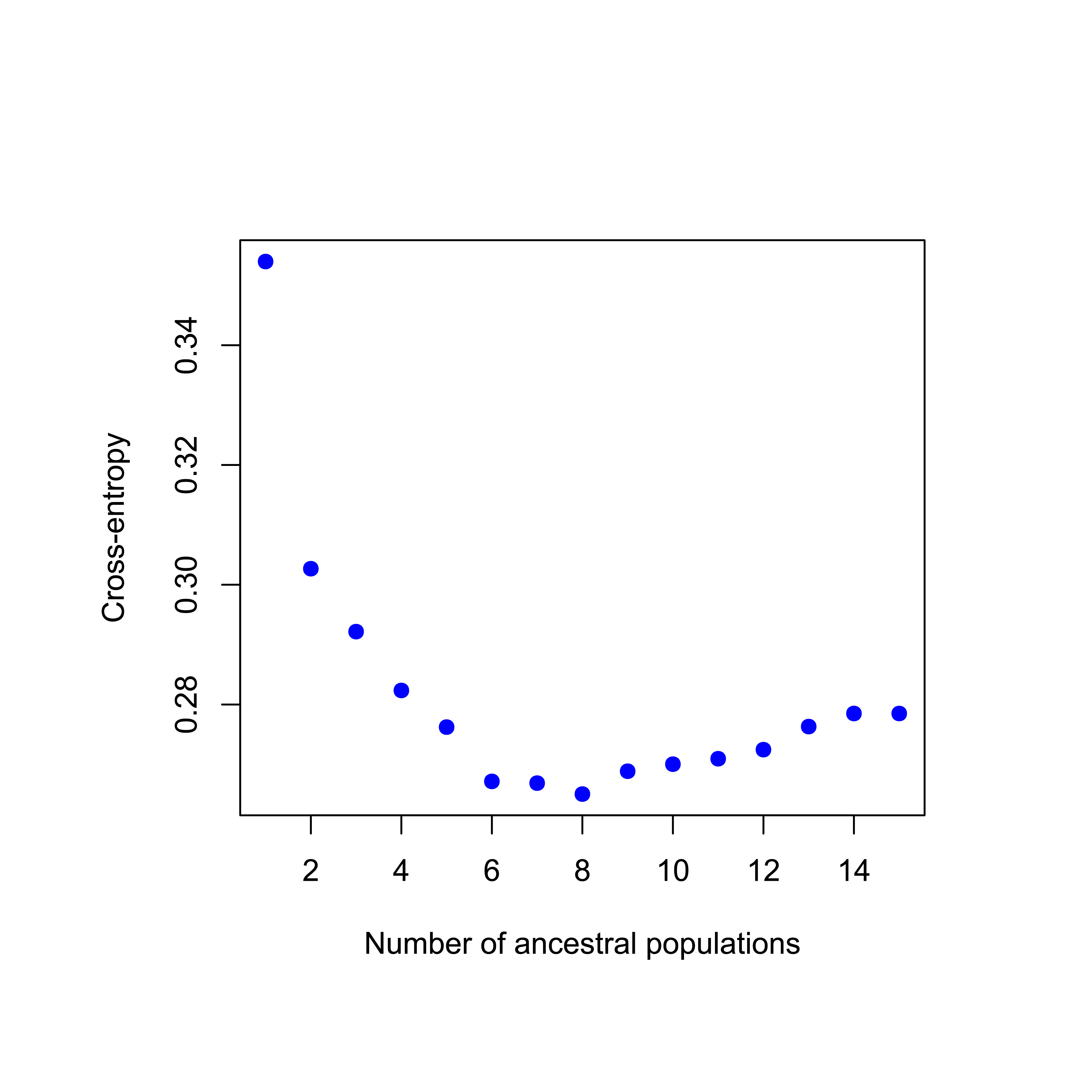

Figure S1. Cross-entropy scores for each number of ancestral populations in the sNMF analysis.

Alt text: A plot depicting data as blue points with number of ancestral populations on the x-axis and cross-entropy scores on the y-axis. Cross-entropy scores decrease until reaching eight ancestral populations and plateau at six ancestral populations. Cross-entropy scores increase after eight ancestral populations.

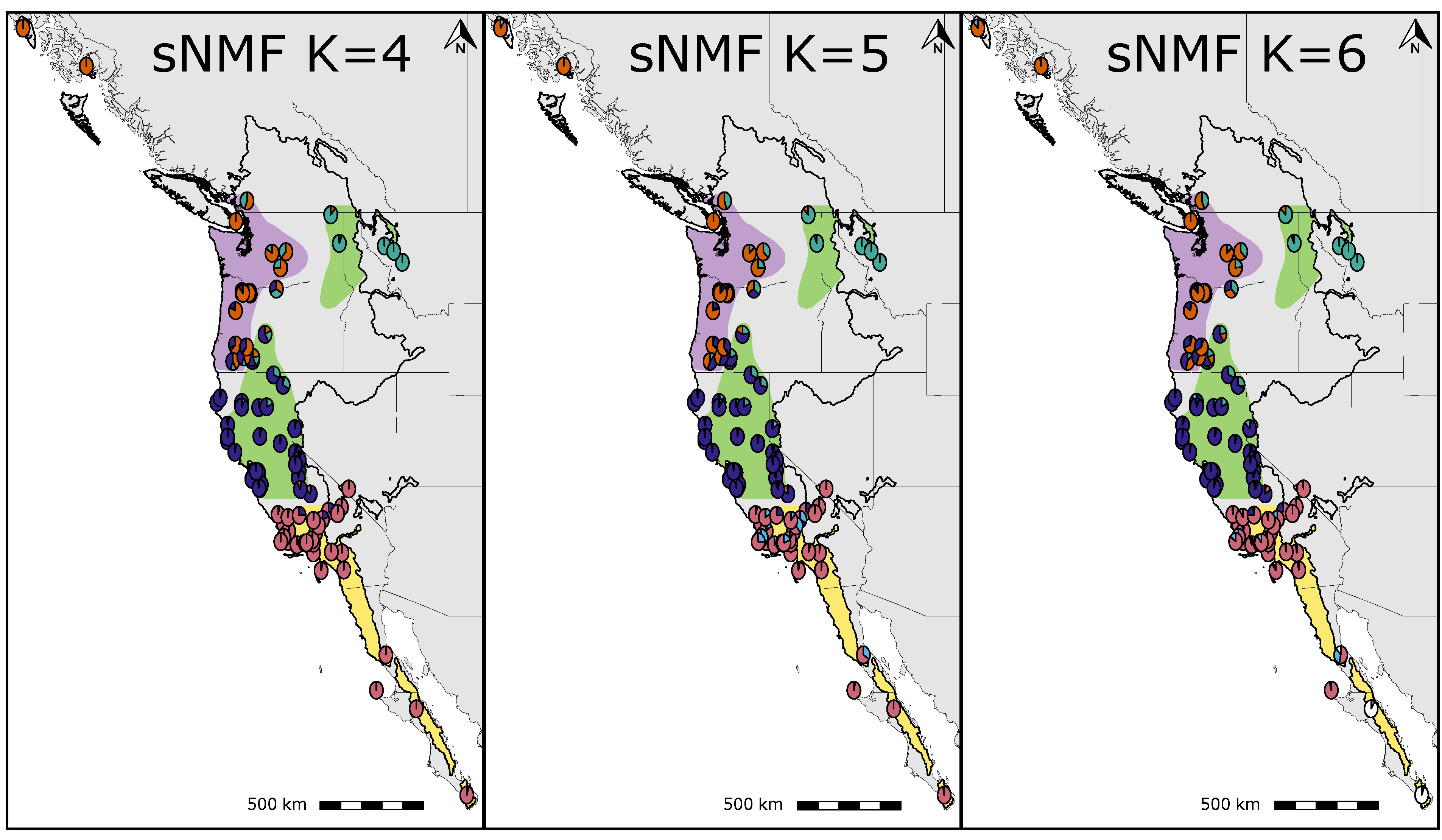

Figure S2. sNMF results for 4 – 6 ancestral populations (K = 4 – 6). Each pie chart represents an individual sample and its ancestry proportions. Colored ranges are based on ranges for *Pseudacris hypochondriaca*, *P. regilla*, and *P. sierra* we approximated from the localities in Jadin et al. (2021). The complete range of *P. regilla sensu lato* is outlined in black.

Alt text: sNMF results displayed as pie charts on maps. K = 4 splits the “coastal” cluster from K = 3 into a northwest cluster and a central California cluster. K = 5 does not form a distinct fifth cluster and instead shows ancestry scattered within the *P. hypochondriaca* samples. K = 6 forms a population cluster for Baja California Sur samples.

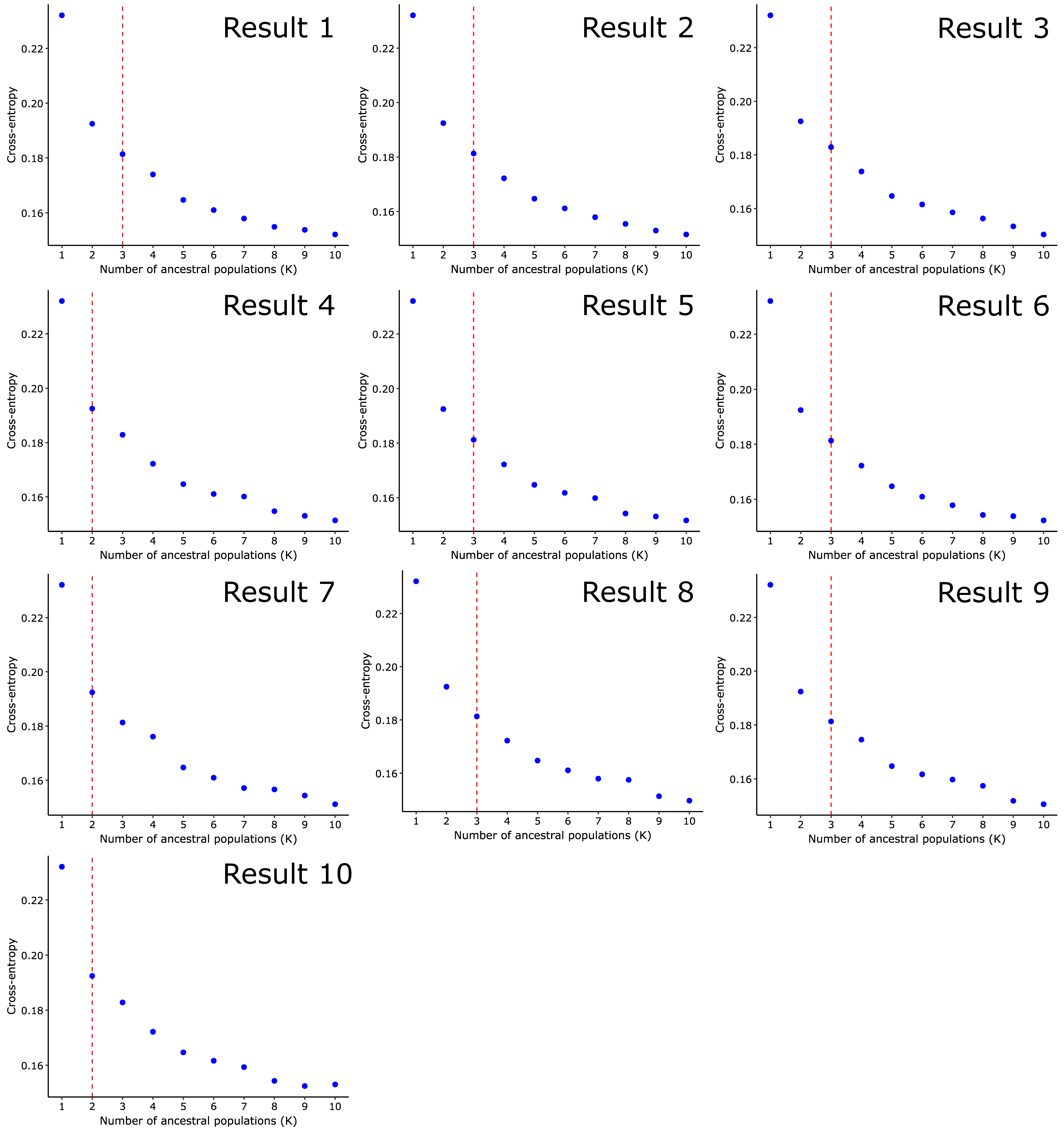

Figure S3. Cross-entropy scores for each number of ancestral populations (K) in the ten TESS3 runs. The dashed red lines represent the “best” value of K for each run.

Alt text: Ten plots depicting data as blue points with number of ancestral populations on the x-axes and cross-entropy scores on the y-axes. Cross-entropy scores decrease as the number of ancestral populations increase. The “best” K is 2 for results 4, 7, and 10. The remaining plots display K = 3 as the “best” number of ancestral populations.

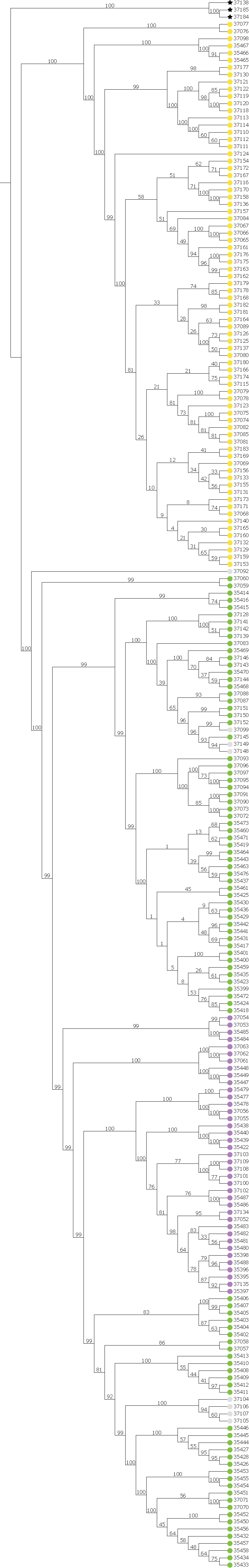

Figure S4. Full SVDQuartets phylogeny. Bootstrap values are included on each branch. Colored circles at the tips of branches represent species assignments based on Jadin et al. (2021).

Alt text: A phylogeny without branch lengths displaying all samples.

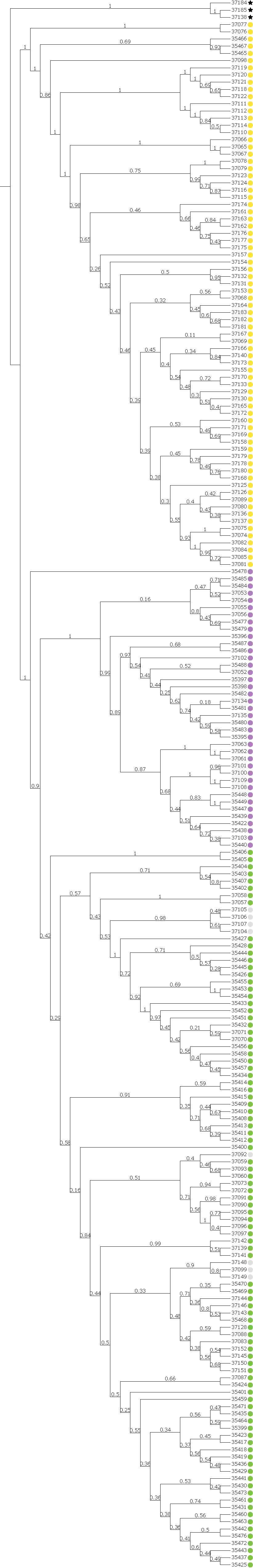

Figure S5. Full ASTRAL phylogeny for the sequence dataset containing at least 98% of individual samples and no more than 5% missingness. Bootstrap values are included on each branch. Colored circles at the tips of branches represent species assignments based on Jadin et al. (2021).

Alt text: A phylogeny without branch lengths displaying all samples.

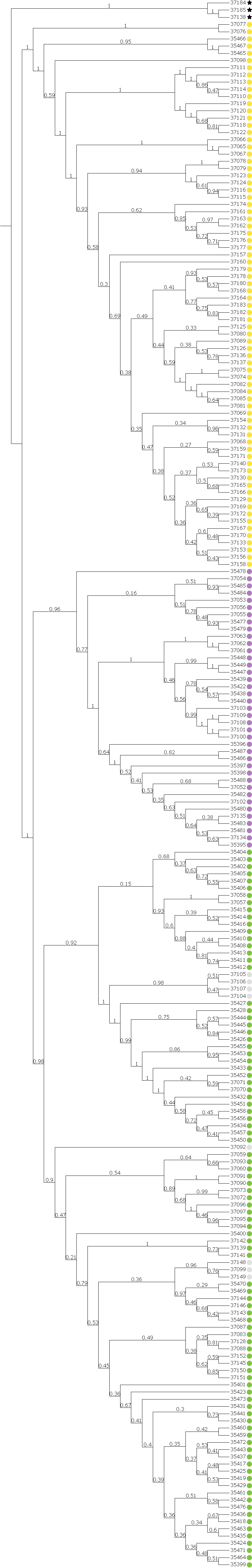

Figure S6. Full ASTRAL phylogeny for the sequence dataset containing at least 98% of individual samples and no more than 10% missingness. Bootstrap values are included on each branch. Colored circles at the tips of branches represent species assignments based on Jadin et al. (2021).

Alt text: A phylogeny without branch lengths displaying all samples.

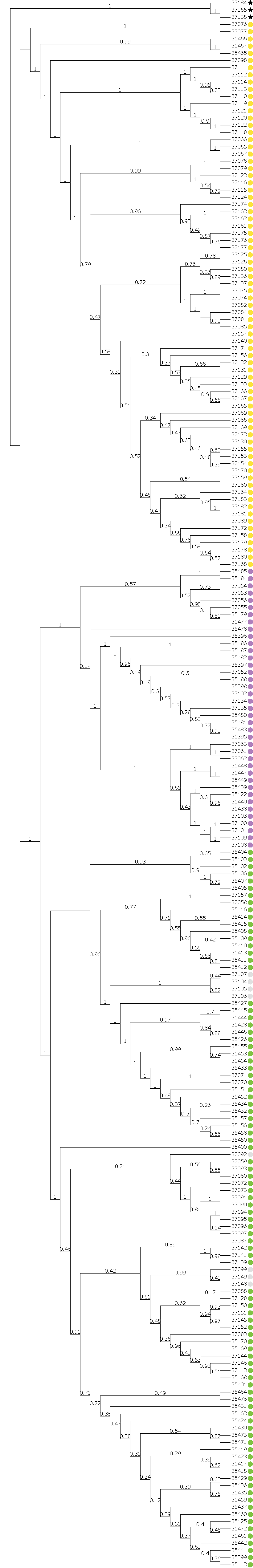

Figure S7. Full ASTRAL phylogeny for the sequence dataset containing at least 92% of individual samples and no more than 5% missingness. Bootstrap values are included on each branch. Colored circles at the tips of branches represent species assignments based on Jadin et al. (2021).

Alt text: A phylogeny without branch lengths displaying all samples.

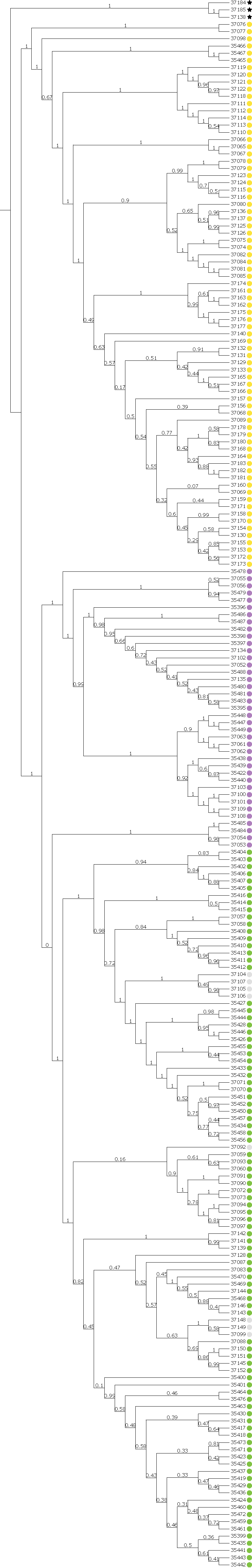

Figure S8. Full ASTRAL phylogeny for the sequence dataset containing at least 92% of individual samples and no more than 10% missingness. Bootstrap values are included on each branch. Colored circles at the tips of branches represent species assignments based on Jadin et al. (2021).

Alt text: A phylogeny without branch lengths displaying all samples.

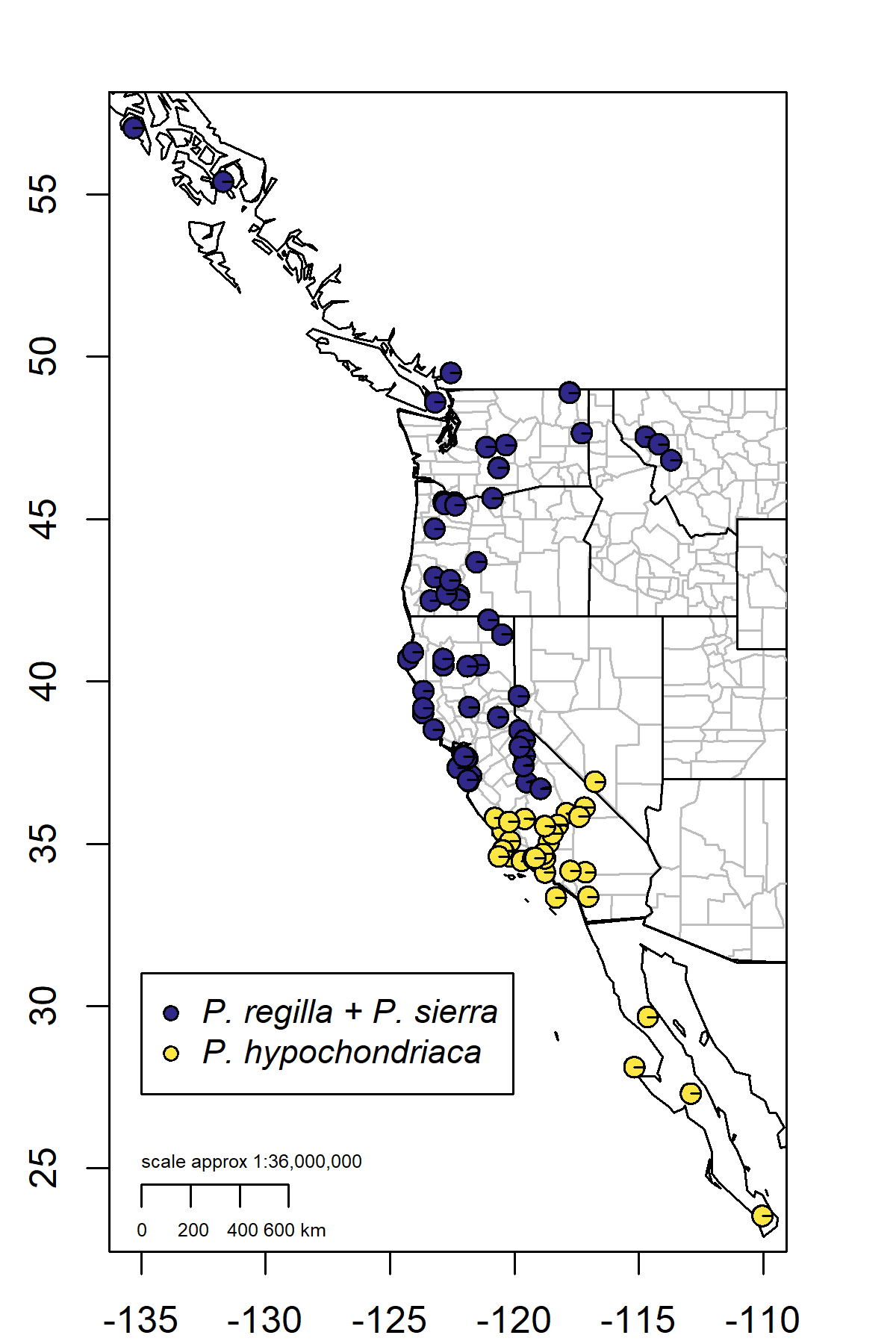

Figure S9. Species coefficients for each sample estimated from the three-layer SuperSOM using the nuclear genomic dataset generated in this study.

Alt text: Species coefficients are displayed as colored pie charts on a map.

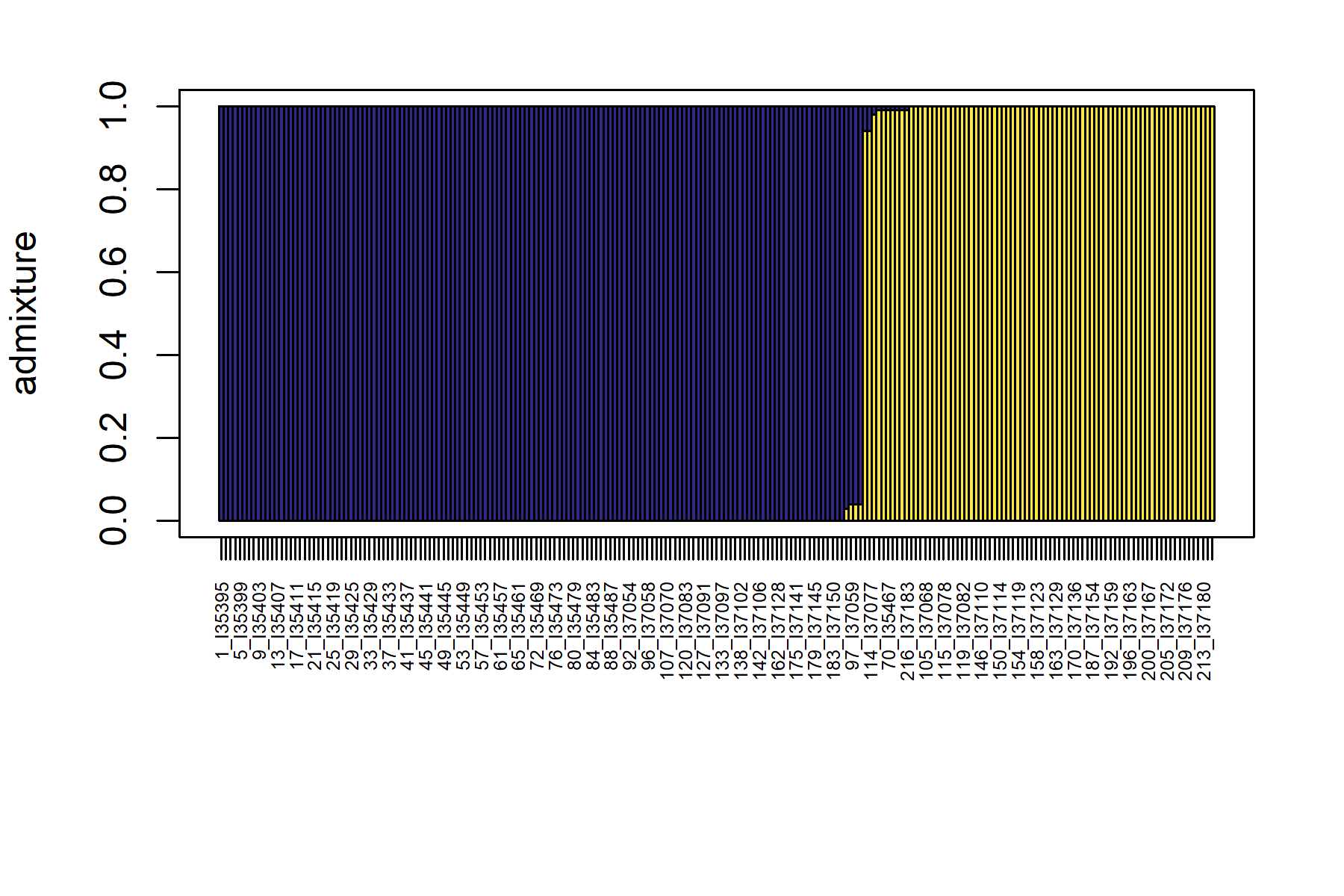

Figure S10. Species coefficients for each sample estimated from the three-layer SuperSOM using the nuclear genomic dataset generated in this study.

Alt text: Species coefficients are displayed as a structure-like colored bar plot with each bar representing a separate sample.

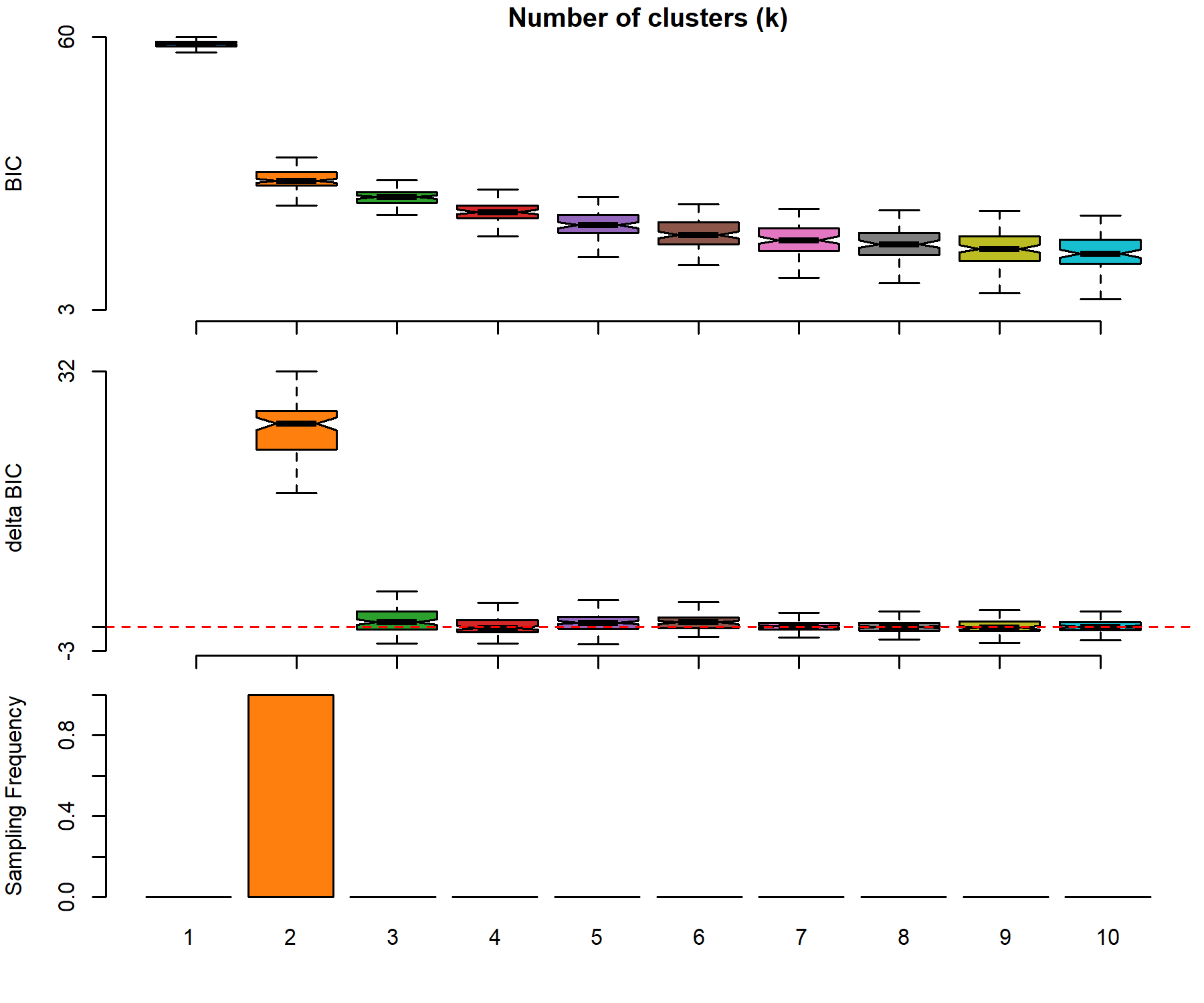

Figure S11. Optimal number of clusters (K) for the four-layer SuperSOM using the dataset from Vélez and Ingram (2025). Bayesian Information Criterion, change in Bayesian Information Criterion, and the sampling frequency for each K of 1-10 are reported.

Alt text: SuperSOM BIC and change in BIC are displayed as box plots. Sampling frequency is displayed as a bar plot.

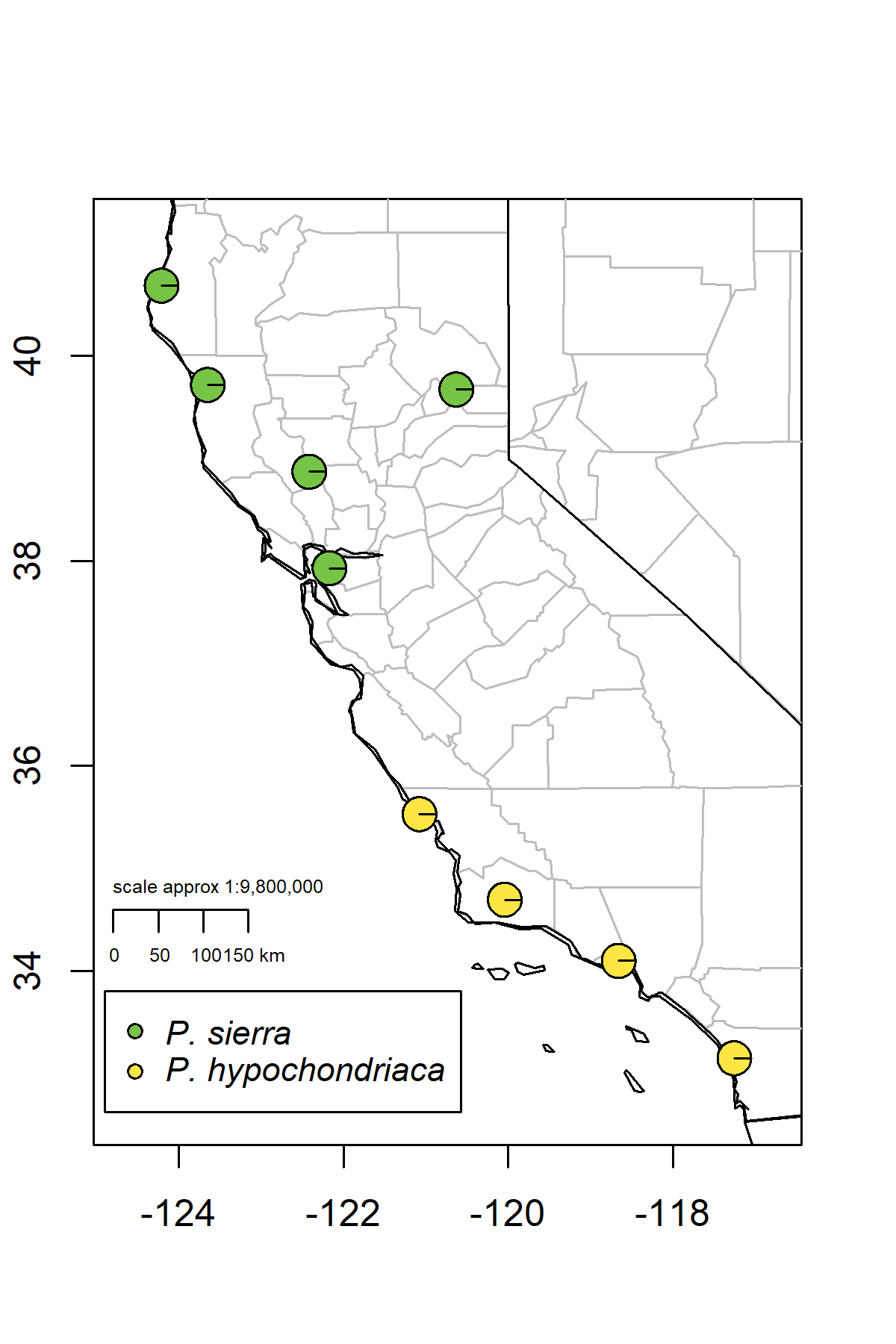

Figure S12. Species coefficients for each sample estimated from the four-layer SuperSOM using the dataset from Vélez and Ingram (2025).

Alt text: Species coefficients are displayed as colored pie charts on a map.

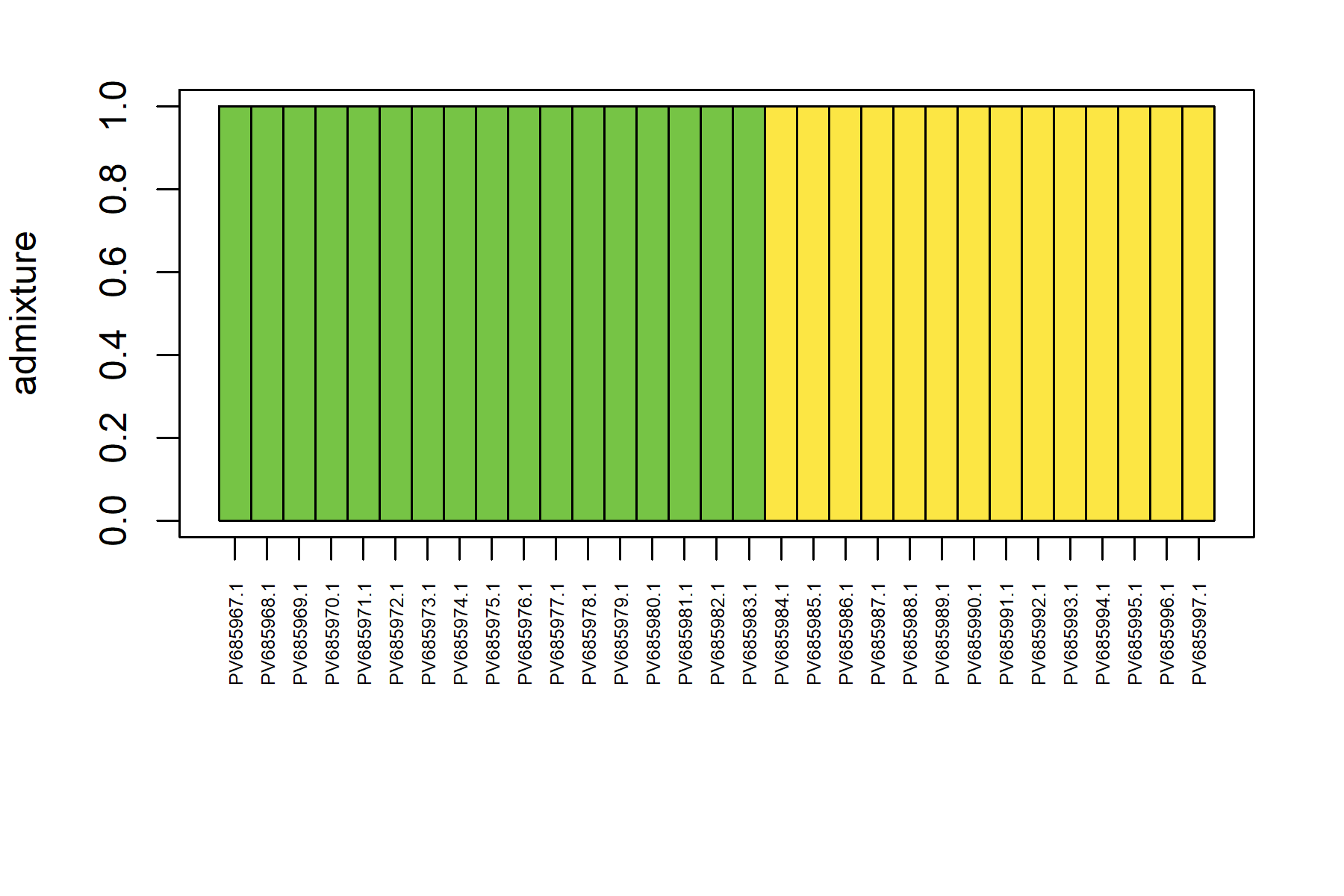

Figure S13. Species coefficients for each sample estimated from the four-layer SuperSOM using the dataset from Vélez and Ingram (2025).

Alt text: Species coefficients are displayed as a structure-like colored bar plot with each bar representing a separate sample.

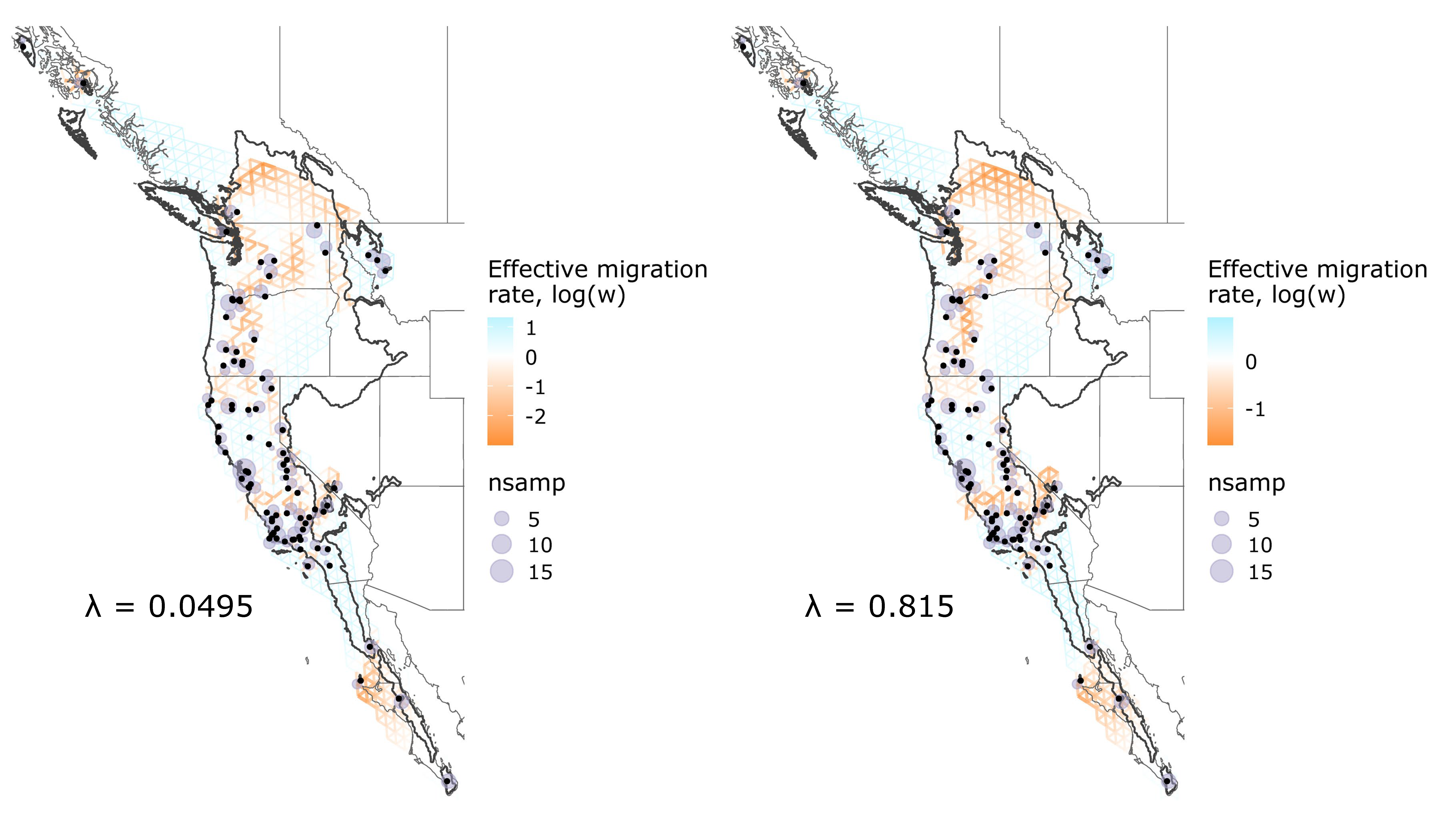

Figure S14. FEEMS results when the smoothing parameter (λ) = 0.0495 (left) and λ = 0.815 (right). Line thickness and color represent the effective migration rate, with orange and thicker lines representing lower effective migration rates, and blue and thinner lines representing higher effective migration rates. The number of samples for each node is represented by the size of the purple circle. Points of the original samples are added as black dots. The range of the *P. regilla sensu lato* complex is outlined in black.

Alt text: Each FEEMS map shows a grid of demes across the landscape, representing effective migration surfaces.
